# Effects of Arginine Vasopressin on Islet Cells in Pancreatic Tissue Slices: Glucose-Dependent Modulation of Alpha and Beta Cell Activity

**DOI:** 10.1101/2025.03.03.641205

**Authors:** Jasmina Kerčmar, Nastja Murko, Lidija Križančić Bombek, Eva Paradiž Leitgeb, Johannes Pfabe, Sandra Postić, Ya-Chi Huang, Andraž Stožer, Dean Korošak, Xaver Kozisek, Monika Perisic, Markus Muttenthaler, Christian W. Gruber, Marjan Slak Rupnik

**Author notes:** Correspondence should be addressed to Marjan Slak Rupnik and Christian W. Gruber. Equal contribution (Jasmina Kerčmar, Nastja Murko, Lidija Križančić Bombek).

## Abstract

Arginine vasopressin (AVP) is well known for regulating fluid volume, osmotic balance, and vascular tone. Its role in the regulation of pancreatic α and β cell function has been reported, yet its effects are not fully understood, particularly regarding its interaction with plasma glucose levels. The osmotic and volume challenges posed by hyper- and hypoglycaemia in diabetes can be a significant complication of effective hormonal regulation of metabolism. In this study, we primarily investigated the effects of AVP and synthetic peptide receptor agonists and antagonists on α and β cells in pancreatic tissue slices using live confocal Ca^2+^ imaging. Our findings demonstrate that AVP exerts glucose-dependent effects on both cell types. At low glucose concentrations, AVP, in combination with physiologically or pharmacologically increased cAMP levels, selectively activated α cells without significantly affecting β cells. In contrast, at higher glucose concentrations and pharmacologically elevated cAMP levels, physiological levels of AVP enhanced β cell activity, leading to increased Ca^2+^ oscillations and insulin release. In both cell types, AVP displayed a bell-shaped concentration dependence, with lower AVP concentrations stimulating hormone release and higher concentrations leading to diminished responses, consistent with inositol trisphosphate receptor (IP_3_R) activation and inactivation properties. Furthermore, our results indicate that AVP acts primarily through V_1b_ receptors in β cells, with no involvement of V_1a_, V_2_ or oxytocin receptors. These findings provide new insights into the modulation of glucose-dependent release of pancreatic hormones by AVP in the context of changed blood osmolality due to hyper- or hypoglycemia in diabetes. Importantly, our results emphasize the potential of targeting AVP signaling pathways as a therapeutic approach in diabetes research, aiming to improve hormone regulation and nutrient homeostasis.

**Highlights:**

- Highly spatio-temporally resolved imaging of islet Ca^2+^ oscillations on pancreatic tissue slices provides an in situ-like model for physiological and pharmacological approaches.
- Physiological glucose stimulation triggers non-linear β cell collective responses that must be taken into account when interpreting single concentration pharmacological experiments.
- In a high cAMP context, AVP acts through V_1b_ receptors on islet α and β cells, exhibiting a bell-shaped dependence driven by the activation-inactivation properties of IP₃ receptors.
- AVP modulates glucose-dependent effects on α and β cells in a physiological concentration range, in the presence of altered blood osmolality or volume due to hyperglycemia, or to the direct effects of hypoglycemia in diabetes.

## Introduction

The most extensively studied physiological roles of arginine vasopressin (AVP; a.k.a. ADH) include the regulation of body fluid volume by promoting water reabsorption in the kidneys, maintenance of osmotic and electrolyte balance, and contribution to cardiovascular homeostasis through regulation of vascular tone and arterial pressure (1, 2). Other actions of AVP, particularly its connection to aspects of metabolic syndrome, are less frequently highlighted (3, 4). AVP has been described to elevate blood glucose levels through two primary mechanisms: enhancing glycogenolysis in the liver and promoting glucagon release from pancreatic islets (5, 6). However, the direct effects of AVP agonists and antagonists on pancreatic β and α cells remain contentious, with ongoing debates in the literature (7–10). This uncertainty is partly because systemic AVP responses can originate from extra-islet targets, making it necessary to examine AVP actions directly within intact pancreatic tissue.

The physiology of pancreatic β cells, which secrete insulin, and α cells, which secrete glucagon, are intricate and remains incompletely understood. The principal hormones released by these cell types are essential for metabolic homeostasis, responding dynamically to nutrient availability and metabolic intermediates present in the plasma (4, 11). These responses to nutrients are further modulated by neurohormonal signals, including AVP. AVP is a nonapeptide synthesized mainly in magnocellular AVP neurons in the supraoptic and paraventricular nuclei of the hypothalamus (12). Its release from the posterior pituitary into the systemic circulation is mainly stimulated by increased plasma osmolality, substantial blood-volume depletion, and diverse neuronal, endocrine or metabolic inputs to the hypothalamus (13). The resulting physiologically active plasma AVP concentration in rodents and humans has been reported to be in the low to mid-picomolar range (14–16). Some studies suggest that circulating AVP and the AVP produced in the pancreas could affect insulin and glucagon secretion via paracrine signaling (17).

One significant factor contributing to the controversies regarding the direct effects of AVP is the glucose concentration used in experimental settings. Insulin and glucagon secretions are reciprocally correlated with plasma glucose concentrations (18). While glucose alone stimulates oscillations in intracellular Ca^2+^ concentration in pancreatic β cells and causes insulin secretion (19), AVP has been reported to act as a positive modulator of glucose-stimulated insulin release (20). Notably, when AVP is administered intraperitoneally together with glucose in fasted mice, there is no significant increase in overall insulin secretory activity compared to glucose alone; however, it results in a marked reduction in blood glucose levels (8). Independently of glucose levels, several in vivo studies across species utilizing intravenous or intramuscular AVP administration have reported increased pancreatic hormone release (6, 13, 20–22). Conversely, other studies have reported no or selective effects on glucagon (23, 24) and insulin release (6, 25), and in vitro experiments in rats have also described AVP-dependent inhibition of insulin release (25). In addition to its effects on insulin secretion, AVP has been reported to enhance proliferation of rodent and human β cells and to protect against cytokine-induced β cells apoptosis (8).

In an organism, AVP exerts its diverse physiological actions by binding to four transmembrane G-protein-coupled receptors (GPCRs): V_1a_, V_1b_, V_2_, as well as oxytocin receptors (26). These receptors differ in their tissue expression, signaling mechanisms, and physiological functions. V_1a_ receptors are primarily expressed in vascular smooth muscle, heart, liver, and central nervous system, whereas V_1b_ receptors are found in the anterior pituitary gland and pancreas. Activation of V_1a_ receptors is associated with vascular contraction, cardiac function (13) and modulation of glucose and lipid metabolism, whereas activation of V_1b_ receptors is involved in the release of adrenocorticotropic hormone (ACTH), glucagon and insulin. Both receptor types couple to the G_q/11_ proteins and activate the phospholipase C (PLC) pathway, leading to increased intracellular Ca^2+^ levels. In contrast, V_2_ receptors preferentially couple to the G_s_ proteins and activate the adenylate cyclase pathway, increasing cyclic AMP (cAMP). This signaling cascade is crucial for water reabsorption in the kidneys, where V_2_ receptors are primarily expressed in the renal collecting ducts and vascular endothelial cells, facilitating the insertion of vesicles containing aquaporin channels into the cell membrane, thereby enhancing water permeability (27).

For pancreatic islets specifically, available transcriptomic datasets indicate that V_1b_ receptor expression is strongly enriched in α cells, whereas V_2_ receptor expression is absent or very low in endocrine islet cells, and β cell V_1b_ receptor expression appears low and remains difficult to resolve conclusively (28). The binding of AVP to either V_1a_ or V_1b_ receptor and activation of PLC leads to intracellular Ca^2+^ release due to inositol trisphosphate (IP_3_) production. Such IP_3_-mediated Ca^2+^ release has been shown to stimulate the secretion of glucagon (13) or insulin in primary (20) and clonal β cell preparations (29). Pharmacological studies have also supported the role of G_q/11_ coupled receptors, as specific V_1a_ or V_1b_ receptor antagonists abolished, and an oxytocin receptor antagonist partially reduced an insulinotropic action of AVP in rodent BRIN BD11 β cells (8). Nevertheless, recent transcriptomic studies support α cells as the dominant V_1b_ receptor-expressing endocrine population, and therefore β cell responses to AVP may reflect a combination of low-level direct receptor-depending effects, indirect intra-islet signaling, and state-dependent modulation of the β cell collective. The potential involvement of V_1a_ receptors in α cells has also not been completely ruled out (8). Studies using genetically modified rodents have provided further evidence for the role of AVP and V1_b_ receptors in islet function. Mice deficient in V_1b_ receptors could not release insulin after AVP stimulation (7, 30), with no difference in fasting blood glucose levels (7). Similarly, glucagon levels were significantly reduced in V_1b_^−/−^ mice (9). On the other hand, Brattleboro rats, which carry a naturally occurring mutation that abolishes AVP production, have been reported to have improved glucose tolerance (31).

It is important to emphasize that AVP does not act primarily as an initiator of hormone release, but rather has a permissive and potentiating role (32). For instance, AVP facilitates corticotropin-releasing hormone (CRH) (33) or histamine-dependent (34) release of adrenocorticotropic hormone (ACTH) from the anterior pituitary, both of which involve G_s_/cAMP-dependent-signaling. β and α cells also utilize signaling pathways involving G_s_-coupled protein receptors, where the permissive and potentiating roles of AVP may be significant (35, 36). In islets, this is particularly relevant because cAMP-elevating conditions may amplify the secretory consequences of Gq-dependent Ca²⁺ signals, although this does not necessarily mean that the upstream AVP-evoked Ca²⁺ response itself is cAMP-dependent. Thus, experiments performed in cAMP-elevated conditions should be interpreted as testing AVP action in a permissive signaling background, rather than as proof of a direct cAMP requirement for AVP receptor activation.

Finally, because the downstream readout of V_1b_ receptor activation involves IP_3_ receptors with well-described activation and inactivation properties (37), such an arrangement could accommodate the broad spectrum of previously observed effects of AVP on the function of α and β cells. However, the relative contributions of direct receptor activation, intra-islet paracrine interactions, and the current dynamic state of the β cell population remain unresolved. On that note, the present study reassessed the effects of AVP and the selection of specific novel AVP receptor agonists and antagonists on pancreatic β and α cells using fresh pancreatic tissue slices combined with live confocal imaging of intracellular Ca^2+^ oscillations. Pancreatic slices provide a complementary preparation that preserves local tissue architecture and intercellular interactions, while also carrying limitations shared with other ex vivo preparations, including the absence of intact circulation and innervation. In this preparation, the majority of AVP stimulatory activity occurs within the known physiological AVP range (1), a range that correlates with the correction of plasma osmolarity through water retention in the distal nephron or oral water intake, and with physiological scenarios related to blood volume depletion.

## Material and methods

### Ethics Statement

The study strictly adhered to all national and European guidelines regarding the care and treatment of laboratory animals. Every effort was made to minimize animal suffering and improve animal welfare. The experimental protocol received approval from the Administration of the Republic of Slovenia for Food Safety, Veterinary and Plant Protection (grant No.: U34401-12/2018/2) and The Ministry of Education, Science and Research, the Republic of Austria (GZ 2022-0.325.009).

### Animals, Tissue Slice Preparation and Indicator Loading

Experiments were conducted 8–25-week-old C57BL/6J mice of both sexes, which were housed in individually ventilated cages (Allentown LLC, USA) at a room temperature 22-24 °C and 45-55% relative humidity with a 12-hour light/dark cycle. The mice had *ad libitum* access to water and a standard chow (Ssniff, Soest, Germany). Acute pancreatic tissue slices were prepared as previously described (38). Briefly, after CO_2_ anesthesia, cervical dislocation and opening of the abdominal cavity, the common bile duct was clamped distally at the major duodenal papilla 1.9% low melting point agarose (Lonza, Basel, Switzerland) was injected proximally. Agarose was dissolved in an extracellular solution (ECS) containing (in mM) 125 NaCl, 26 NaHCO_3_, 6 glucose, 6 lactic acid, 3 myo-inositol, 2.5 KCl, 2 Na-pyruvate, 2 CaCl_2_, 1.25 NaH_2_PO_4_, 1 MgCl_2_, 0.5 ascorbic acid and kept in a prewarmed water bath at 40 °C. Following the agarose injection, the pancreatic tissue was immediately cooled with ice-cold ECS, extracted from the abdominal cavity, and placed into a large Petri dish containing ice-cold ECS. Next, 3-5 mm^3^ large tissue cubes were cut from the pancreas, cleared from the connective tissue, and embedded into agarose. The embedded tissue blocks were sectioned into 140 µm thick slices using a vibratome (VT1000S, Leica Microsystems, Wetzlar, Germany). The tissue slices were stored at room temperature in HEPES-buffered saline containing 6 mM glucose (HBS, consisting of (in mM) 150 NaCl, 10 HEPES, 6 glucose, 5 KCl, 2 CaCl_2_, 1 MgCl_2_; titrated to pH=7.4 with 1 M NaOH). For Ca^2+^ indicator loading, the slices were incubated for 50 minutes in 3.33 ml of HEPES-buffered saline containing 6 mM glucose, 3.75 µl dimethylsulfoxide, 1.25 µl Pluronic F-127, and 6 µg Calbryte 520 AM (AAT Bioquest, Pleasanton, CA, USA). Unless specified otherwise, all chemicals were obtained from Sigma Aldrich (St. Louis, MO, USA).

### Immunofluorescence and RNAscope

Fresh-frozen samples were sectioned at 20 µm using a cryostat (Leica CM1950). Tissue fixation was performed in cold 4% paraformaldehyde (PFA) for 1 h. Sections were blocked with 5% BSA and 0.3% Triton X-100 in PBS for 1 h at room temperature (RT). Incubation with the somatostatin primary antibody (Sigma-Aldrich, SAB4502861) was carried out overnight at 4 °C. The secondary antibody (Invitrogen, A-21245) was incubated for 2 h at RT. To assess tissue morphology, DAPI staining was performed at a concentration of 1 µg/mL.

RNAscope was performed using the RNAscope™ Multiplex Fluorescent Reagent Kit v2 (ACD Bio-Techne, 323136) according to the manufacturer’s recommended conditions and supplied reagents, with minor modifications (39). Fresh-frozen samples were sectioned at 20 µm and dried inside a cryostat (Leica CM1950) for 2 h at −26 °C. When not used immediately, slides were stored at −80 °C according to the manufacturer’s instructions. Protease IV digestion was prolonged by an additional 10 min, and the number of wash steps was increased to three washes of 5 min each. The RNAscope™ probes used were Probe Mm-Avpr1b C1 (ACD Bio-Techne, 480141) and Probe Mm-Gcg C2 (ACD Bio-Techne, 400601-C2). For probe detection, TSA Vivid dyes 520 (ACD Bio-Techne, 323271) and 570 (ACD Bio-Techne, 323272) were used at a 1:1500 dilution, respectively. Probe visualization was acquired using a confocal microscope (Olympus IXplore SpinSR) equipped with a Hamamatsu ORCA-Fusion 2 camera. Overview images were acquired using a 20× dry objective, and islet images were acquired using a 60× silicone oil immersion objective (NA 1.3).

The cell segmentation of islet images was based on the fluorescent DAPI signal. Briefly, the 2D Gaussian filter of a kernel size of 20 was applied over the DAPI channel and the background signal was removed using Otsu-thresholding (40). Next, the local maxima was found defining the nuclei center. As a last step, watershed (41) segmentation was used with a maximum cell radius of 150 pixels (∼15 µm) using the detected peaks. RNAscope signal was counted by applying a maximum entropy thresholding on the respective fluorescence data. Next, we detected dots by using a Laplacian of Gaussian (42) approach with a minimum sigma of 0.75 and a maximum sigma of 2. Detected dots were mapped back to the DAPI segmentation data. Each cell with more than 2 dots was counted as positive.

### Compound Synthesis, Purification, and Quality Control

VP analogs [(D-Leu)^2^,Ile^3^,Thr^4^]-VP, d[Cha^4^,Dab^8^]-VP, and d[(CH_2_)_5_^1^,Tyr(Me)^2^,Dab^5^,Tyr^9^]-VP were prepared following established procedures *via* Fmoc-SPPS (9-fluorenylmethyloxycarbonyl solid-phase peptide synthesis) and standard purification and quality control methods (43) (Fig. 5, Suppl. Fig. 1, Suppl. Table 1). Shortly, peptides were obtained through manual synthesis on a Rink amide aminomethyl resin followed by TFA cleavage, oxidative folding in a 0.1 M ammonium bicarbonate buffer (pH 8.2, 24 h), and purification *via* reversed phase-high performance liquid chromatography (RP-HPLC) on a Waters Auto Purification HPLC-UV system (Kromasil Classic C_18_ 21.2 x 250 mm, 300 Å, 10 μm) at a solvent flow of 20 mL/min and a linear gradient of 5-45% solvent B in solvent A (A: ddH_2_O + 0.1 % TFA, B: ACN + 0.08 % TFA) in 50 min. Reactions were monitored by analytic RP-HPLC-mass spectrometry (RP-HPLC-MS) on a Thermo Scientific Dionex Ultimate 3000 system equipped with a Waters XSelect CSH UPLC C_18_ XP column (3.0 x 75 mm, 130 Å, 2.5 µm), UV detection (measurement at 214 nm and 280 nm), and Thermo Scientific MSQ Plus ESI-MS unit (positive ionization mode). Compound purity was determined through analytical RP-HPLC peak integration at 214 nm, and compound identity was verified through direct injection MS (Supplementary Information). A Thermo Fisher Scientific Vanquish Horizon UHPLC system using a designated column (Kromasil Classic C_18_ 2.1 x 100 mm, 100 Å, 5 µm) was used to determine the product concentration based on peptide absorption at 214 nm, and a comparison of absorption areas to peptide standards with known concentration (44).

### Pharmacological Characterization

HEK293 cells stably expressing the EGFP-tagged receptor of interest (human oxytocin, V_1a_, or V_1b_ receptor) were cultured. Radioligand displacement was performed in duplicates of cell membranes (6–20 µg) incubated with the radioactive agonist ^3^H-OT or ^3^H-VP and the competing peptide according to previously published protocols (45–47). The used radioligand concentrations refer to the K_d_ of each receptor subtype which was obtained from saturation binding experiments (0.65 nM for OTR, 0.21 nM for V_1a_R and 0.14 nM for V_1b_R). Nonspecific binding was determined by addition of 10 μM OT or VP. After one hour, the reaction was stopped by filtering through glass fiber filters, which were subsequently used to determine the retained radioactivity measured by liquid scintillation.

Activation of G_q/11_-signalling was measured by IP_1_ quantification using the homogeneous time-resolved fluorescence IP-One assay kit (Revvity) according to the manufacturer’s recommendations. Briefly, stable cell lines were seeded at a density of 10,000 cells per well and incubated for 2 days. At the time of the assay, the cells were equilibrated in the provided stimulation buffer for 15 min prior to the addition of the peptide ligands. After 1 h at 37°C, the stimulation was stopped by adding labeled immunoreagents. The mixtures were then incubated for a further hour at room temperature before the fluorescence ratio 665/620 nm was measured on a FlexStation 3 plate reader (Molecular Devices, USA) at an excitation wavelength of 340 nm.

Data analysis of the pharmacological characterization was performed with GraphPad Prism (GraphPad Software, USA) (Fig. 5, Table 1). The potency (EC_50_) and maximum efficacy (E_max_) values were generated by fitting the obtained data to three-parameter nonlinear regression curves with a bottom constrained to zero. Graphs were normalized to the highest response by the control, i.e. oxytocin or vasopressin. For radioligand displacement IC_50_ values were calculated by a three-parameter logistic Hill equation and then further used to approximate *K_i_*values according to Cheng and Prusoff (48). The normalization refers to the specific binding of the radioligand in the absence of peptides as a maximum with an average of 4.2 pmol/mg for OTR, 2.6 pmol/mg for V_1a_R and 2.9 pmol/mg for V_1b_R. All data regarding pharmacological characterization were presented as mean ± SD of at least three independent experiments unless otherwise stated.

**Table 1:** Affinity, potency and efficacy of [(D-Leu)^2^,Ile^3^,Thr^4^]-AVP, d[Cha^4^,Dab^8^]-AVP, and d[(CH) ^1^,Tyr(Me)^2^,Dab^5^,Tyr^9^]-AVP. Affinity (K) and potency (EC) data are indicated as mean ± SD (nM, n = 3); K_i_ values were calculated from IC_50_ values according to Cheng and Prusoff, assuming K_d_ values of 0.65 nM for OTR, 0.21 nM for V_1a_R and 0.14 nM for V_1b_R. Efficacy (E_max_) data is normalized to the maximum IP_1_ formation by the control. ^#^controls were OT at the OTR or AVP at the V_1a_R and V_1b_R; n.d., not determined.

|  | K<sub>i</sub> ± SD (nM) |  |  | EC<sub>50</sub> ± SD (nM) | E<sub>max</sub> ± SD (%) | pA<sub>2</sub> |
| --- | --- | --- | --- | --- | --- | --- |
|  | OTR | V<sub>1a</sub>R | V<sub>1b</sub>R | V<sub>1b</sub>R | V<sub>1b</sub>R | V<sub>1a</sub>R |
| OT | 0.5 ± 0.2 | 18 ± 5 | 588 ± 19 | n.d. | n.d. | - |
| AVP | 8 ± 5 | 0.2 ± 0.1 | 0.3 ± 0.1 | 15 ± 2 | 97 ± 4 | - |
| [(D-Leu)<sup>2</sup>,Ile<sup>3</sup>,Thr<sup>4</sup>]-VP | 209 ± 36 | 5 ± 1 | 70 ± 7 | 361 ± 176 | 79 ± 8 | n.d. |
| d[Cha<sup>4</sup>,Dab<sup>8</sup>]-VP | 54 ± 19 | 23 ± 5 | 0.2 ± 0.02 | 130 ± 52 | 104 ± 13 | n.d. |
| d[(CH<sub>2</sub>)<sub>5</sub><sup>1</sup>,Tyr(Me)<sup>2</sup>,Dab<sup>5</sup>,Tyr<sup>9</sup>]-VP | 2300 ± 190 | 66 ± 15 | >10,000 | - | - | 5.27 |

Cha = cyclohexylalanine; (CH<sub>2</sub>)<sub>5</sub> = β-mercapto-β,β-cyclopentamethylenepropionic acid; d = deaminated N-terminus; Dab = 2,4-diaminobutyric acid.

### Calcium Imaging

Imaging was performed with upright confocal microscope Leica TCS SP5 AOBS Tandem II (20x HCX APO L water immersion objective, NA 1.0) and inverted confocal microscope Leica TCS SP5 DMI6000 CS (20x HC PL APO water/oil immersion objective, NA 0.7). The Ca^2+^ indicator Calbryte 520 was excited using a 488 nm argon laser, and the emitted fluorescence was detected by Leica HyD hybrid detectors in photon-counting mode (Leica Microsystems, Wetzlar, Germany) in the range of 500-700 nm, as previously described (38). The laser power was adjusted to maintain a satisfactory balance between photobleaching and signal-to-noise ratio. The optical imaging thickness was set to nearly 5 μm to prevent recording from multiple cell layers. Time series were acquired with a frequency of 20 Hz and a resolution of 256 x 256 pixels and pixels size of approximately 1 mm^2^.

### Stimulation Protocol

Before imaging, the stained slices were kept in a substimulatory glucose concentration (HEPES with 6 mM glucose) at room temperature. For Ca^2+^ imaging slices were transferred into the recording chamber on the microscope stage, perfused continuously with carbonated ECS containing 6 mM glucose at 37 °C. After recording basal Ca^2+^ signals in 6 mM glucose, the perfusion was exchanged with stimulatory ECS containing 8 (or 9 mM as indicated) glucose for 15-20 minutes. When a stable second phase plateau response with fast Ca^2+^ oscillations was achieved (49), AVP or AVP receptor agonist/antagonist was added to the perfusion for 10-15 minutes. Following AVP/agonist/antagonist testing, the perfusion was again changed to ECS with either 8 or 9 mM glucose for wash-out period of about 15-20 minutes and then to the substimulatory ECS with 6 mM glucose until the cessation of fast Ca^2+^ oscillations. For AVP concentration-dependent responses, islets were stimulated with 8 mM glucose and 500 nM forskolin until the plateau phase has been achieved. A version of this experiment has been performed with 2.5 nM epinephrine. In both cases, a series of AVP concentrations were applied, each of which was static incubated in the recording chamber for 15 minutes before the next concentration was applied.

### Hormone measurements

For acute insulin release in response to glucose, mouse pancreatic tissue slices were preincubated (1 h) in ESC buffer containing 6 mM glucose at 37°C. The sample for the insulin release from the slices was then collected in ESC with 6 mM glucose for 5 min (basal), after which the solution was replaced with stimulatory glucose in ESC 8 mM glucose or ESC 8 mM glucose plus 10 nM AVP and incubated for 15 min. Each solution replacement step involved washing the dish containing the slices at least 3 times to ensure no remaining insulin being present. Insulin concentration was determined using the Insulin Ultra-Sensitive assay kit from Cisbio (Bagnols-sur-Ceze, France).

Dynamic insulin secretion from pancreatic tissue slices was assessed using an in-house built perifusion system with automated tray handling (Lambda Omnicoll, Lambda Laboratory Instruments). Two perifusion columns were connected to the system, each consisting of three chambers positioned vertically. Three pancreatic tissue slices were placed on perforated slice supports (Biorep), one slice per chamber. To equilibrate the slices to 37 °C and wash out accumulated enzymes and hormones, a 60-min pre-perifusion step was performed using a BSA-HEPES–based solution containing 6 mM glucose and soybean trypsin inhibitor (STI) at a flow rate of 140 µL min⁻¹. Subsequently, the experimental perifusion protocol was applied at the same flow rate. Samples were collected at 5-min intervals into 1.5 mL tubes and immediately placed on ice. All samples were stored at −80 °C until insulin or glucagon concentrations were determined using the corresponding HTRF assay kits (Revvity, 62IN1PEG and 62CGLPEG, respectively).

The stimulation solution contained (in mM): NaCl 125; KCl 2.5; HEPES 10; NaHCO₃ 10; Na₂HPO₄ 1.25; sodium pyruvate 2; ascorbic acid 0.25; myo-inositol 3; lactate 6; MgCl₂ 1; CaCl₂ 2; forskolin 0.0005; BSA 0.1% (w/v); and soybean trypsin inhibitor 0.025% (w/v). Glucose (6 or 8 mM) and AVP (1 pM, 100 pM, or 10 nM) were added according to the experimental protocol. All chemicals were purchased from Sigma-Aldrich (St. Louis, MO, USA), unless otherwise stated.

Insulin and glucagon concentrations measured in perifusate samples were normalized to the total hormone content of the tissue slices from the corresponding perifusion column. At the end of each experiment, the three pancreatic tissue slices from each perifusion column were collected and pooled for lysis in acid ethanol to extract intracellular hormone content. Insulin and glucagon concentrations in the lysates were determined using the same assay as for perifusate samples. Hormone secretion during each stimulation period was calculated as cumulative hormone release by summing hormone concentrations measured in all collected fractions during the respective stimulation period and normalizing the obtained value to the total hormone content of the corresponding lysate. To reduce heteroscedasticity, normalized hormone secretion values were log-transformed before statistical analysis and visualization. Outliers were identified after log transformation using a predefined 1.5×IQR rule within each experimental condition and excluded before statistical testing. Normalized and log transformed values from individual experiments were compared across replicates and presented as median ± IQR.

### Data Analysis

The analysis of Ca^2+^ events has been performed as previously described (49). Briefly, the movies were processed using a custom Python script to detect ROIs corresponding to individual cells automatically. Cells have been identified based on their typical activity pattern and intraislet localization as described before (50, 51). β cells have been inactive at non-stimulatory glucose concentration (6 mM) and activated in a biphasic manner when stimulated with either 8 or 9 mM glucose. In contrast, α cells were still active at 6 mM glucose, progressively inhibited in 8 mM glucose, and further stimulated during epinephrine stimulation. Smooth muscle cells were localized around vascular structures in the slices and were used as a readout of V_1a_ receptor activity. All other cells outside the histologically identifiable islet were discarded from further analysis. In the next step, Ca^2+^ events from each ROI were automatically distilled and annotated (49).

For this study, altogether 172 pancreatic tissue slices have been imaged altogether. In addition to the frequency of events per roi per minute (epmpr), for each detected Ca^2+^ event, we measured the height, duration at half of the amplitude (halfwidth) and area under the curve (AUC) (Fig. 2E). Halfwidths between 0.01 to 100 seconds were collected and were pooled together for the global analysis. For the analysis within individual islets, the parameters from each ROI were checked for normality and then parametric analysis (ANOVA/Tukey HSD post hoc) or non-parametric analysis (Kruskal-Wallis/Dunńs test post hoc) has been performed. A similar strategy has been used when we compared several islets exposed to the same treatment. The rois with less than 5 events in the whole length of recording were eliminated from the analysis. The p-value below 0.05 has been taken as statistically significant.

## Results

### V_1b_ receptor transcript expression is enriched in α cells but not restricted to α cells

Previous transcriptomic analyses of purified pancreatic endocrine cells identified V_1b_ receptor expression as strongly enriched in α cells, whereas expression in β cells was reported to be low or absent (28). These data provided an important reference point for interpreting AVP effects in the islet, but they do not fully resolve receptor distribution within the preserved tissue context. We therefore performed RNAscope in situ hybridization in mouse pancreatic sections to assess the localization of V_1b_ receptor transcripts directly in intact pancreatic tissue (Fig. 1).

**Figure 1.**
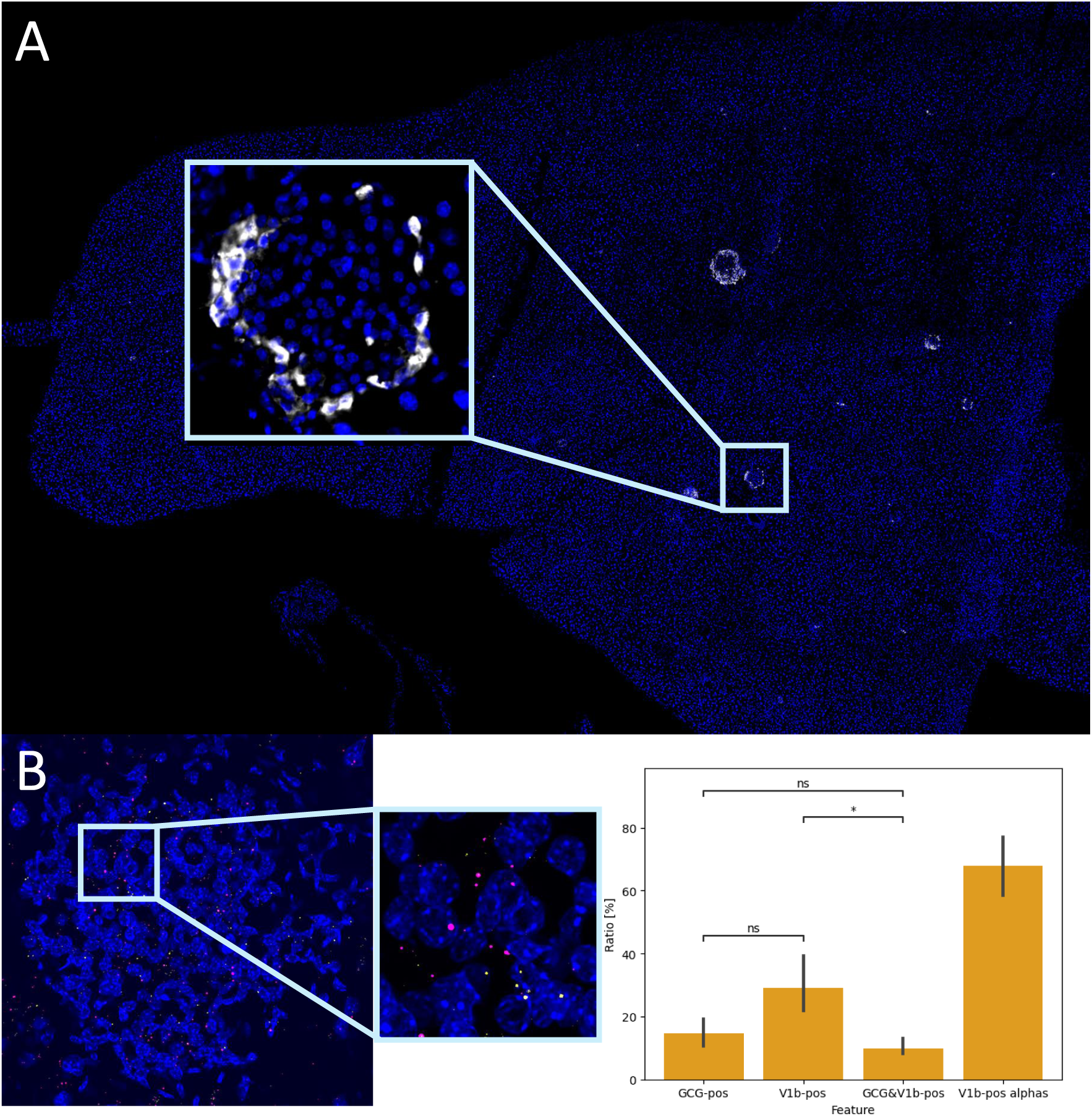
V_1b_ receptor transcript expression in pancreatic islets is enriched in α cells but not restricted to α cells. (A) Mouse pancreatic section showing immunostaining for somatostatin and DAPI, with the islet region shown at higher magnification. (B) RNAscope in situ hybridization of the same islet region. Magenta puncta indicate V_1b_ receptor / Avpr1b transcripts and yellow puncta indicate Gcg transcripts. The magnified inset shows V_1b_ receptor transcript signal in glucagon-positive α cells as well as in other glucagon-negative islet cells. C) Quantification of RNAscope signals showing the relative abundance of glucagon-positive cells, V_1b_ receptor-positive cells, double-positive glucagon/V_1b_ receptor transcript-expressing cells, and the fraction of α cells positive for V_1b_ receptor transcripts. In this dataset, 67.9 ± 7.5% of α cells were V_1b_ receptor-positive, whereas only approximately one third of V_1b_ receptor-positive cells/signals colocalized with glucagon-positive α cells.

Consistent with previous transcriptomic data, V_1b_ receptor transcripts were readily detected in glucagon transcript-positive α cells. However, the signal was not restricted exclusively to glucagon-positive cells, indicating a broader intra-islet expression pattern than would be expected from a strictly α cell-specific marker (Fig. 1A). Quantification confirmed that 67.9 ± 7.5% of α cells were V_1b_ receptor-positive, confirming substantial V_1b_ receptor expression in α cells (Fig. 1B). At the same time, comparison of the V_1b_-positive and glucagon-positive populations indicated that only approximately one third of the V_1b_ receptor-positive signal/cells colocalized with α cells. Thus, V_1b_ receptor expression in the islet is clearly not restricted to α cells.

Together, these data place the functional AVP experiments in the context of previous transcriptomic studies. They confirm that α cells represent an important V_1b_ receptor-positive endocrine population, but also indicate that V_1b_ receptor transcript expression is broader than expected from an exclusively α cell-restricted model. This supports a more cautious interpretation of AVP effects in intact tissue, where β cell modulation may arise from a combination of direct V_1b_ receptor-dependent effects in non-α cells, intra-islet paracrine interactions, and state-dependent collective β-cell dynamics.

### The glucose-dependence of AVP modulation of α and β cell activity

Having established that V_1b_ receptor transcript expression in intact pancreatic islets is not restricted to α cells, we next examined how AVP modulates α and β cell Ca²⁺ activity under different glucose conditions. Since previous work has shown that both AVP responsiveness and islet hormone secretion depend strongly on the metabolic state of the islet, we first tested AVP effects at different glucose concentrations using live Ca²⁺ imaging in acute pancreatic tissue slices.

We have fully reproduced the previously reported glucose-dependent effects of AVP on both α and β cells (Fig. 2). It is crucial to emphasize that at the substimulatory glucose, neither forskolin, an activator of cAMP production alone nor in combination with AVP could significantly activate β cells (Fig. 2BF). On the other hand, forskolin alone or combined with AVP, activated α cells (Fig. 2DG). However, subsequent increase to stimulatory glucose in the same islet activated β cells (Fig. 2B) and progressively reduced the activity of α cells (Fig. 2D). The activation of α cells with AVP did not have a significant effect on β cells (Fig. 2BD).

**Figure 2.**
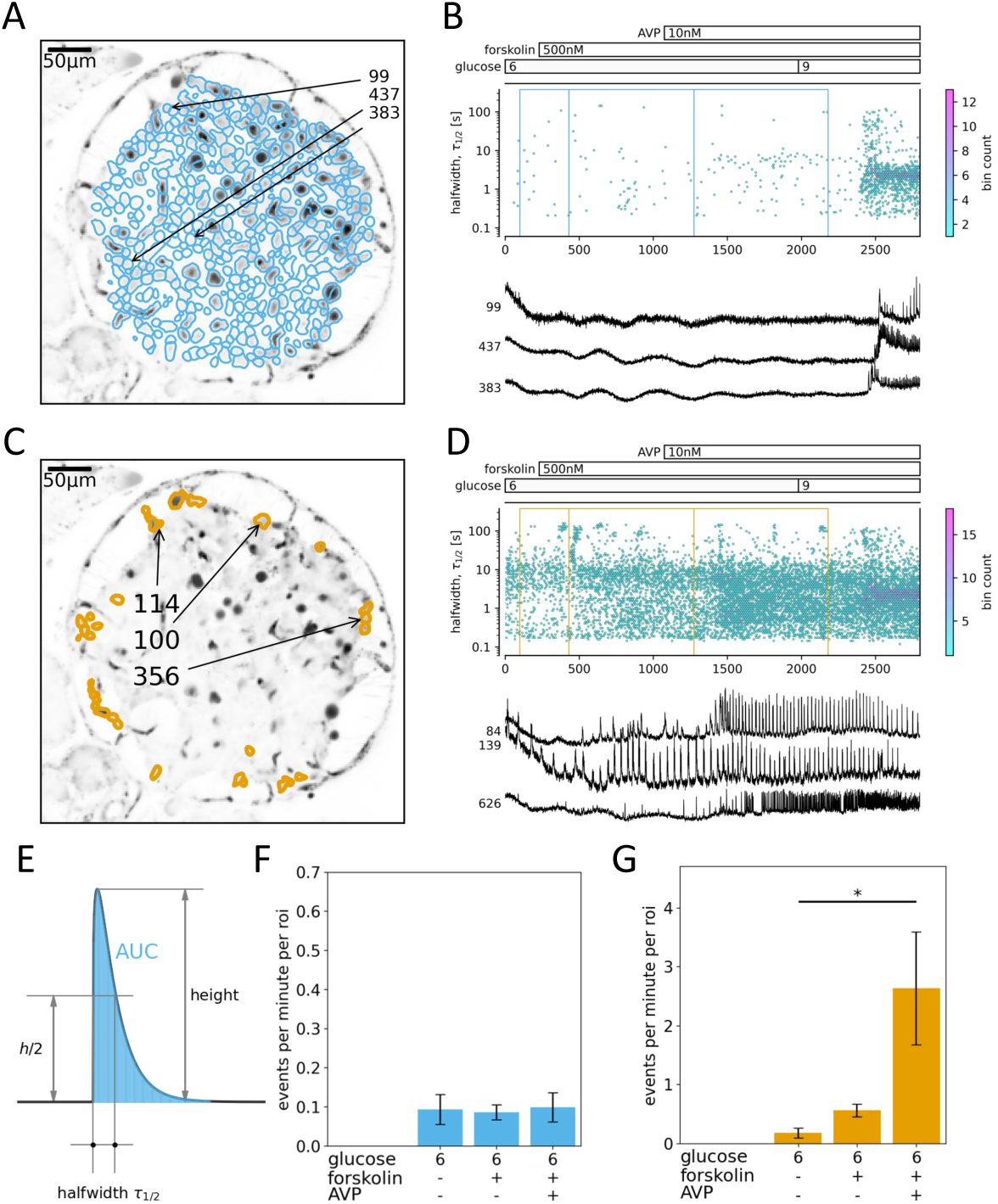
The effect of AVP at the substimulatory glucose on β (A, B, F) and α (C, D, G) cells. (A, C) Regions of interest (ROIs) were obtained with our analytical pipeline. Indicated are the ROIs whose filtered traces correlate best with the average trace for the whole islet. (B, D) (top panel) Time distribution of all measured eventś halfwidths at their peak times. Stimulation protocol is indicated above. The color bar indicates the bin count in each time/halfwidth point. The dashed-line rectangles indicate the 3 time periods where events per minute per ROI parameter was sampled for panels F and G. (bottom panel) Representative traces from ROIs indicated in A and C. (E) A scheme showing how the height, duration at half amplitude (halfwidth) and area under the curve (AUC) were measured for each Ca^2+^ event detected. (F, G) Events per minute per ROI were normally distributed, we used one-way ANOVA and Tukey post-hoc test. The data are presented as mean +/− SD. The significant differences in AVP treatment at the substimulatory glucose were found in α cells only (*p ˂ 0.05).

As previously observed, the application of forskolin to islets after stimulation with physiological levels of glucose (8 or 9 mM), significantly increased the activity of β cells (Fig. 3B). Forskolin, even at low concentration used here (500 nM), increased the frequency of Ca^2+^ oscillations in all islets tested at 8 mM glucose (Fig. 3BE). Application of 10 nM AVP further increased the oscillation frequency, which was often associated with a visible decrease in the halfwidth of the events (Fig. 3B) and in the overall AUC, a parameter that reflects both changes in oscillation frequency and duration (Fig. 3H), although this was not significantly different after pooling the data (Fig. 3GH). It is worth noting that the AUC values varied significantly between different islets, ranging from an increase in overall activity after AVP application to no change or even a decrease in AUC.

**Figure 3.**
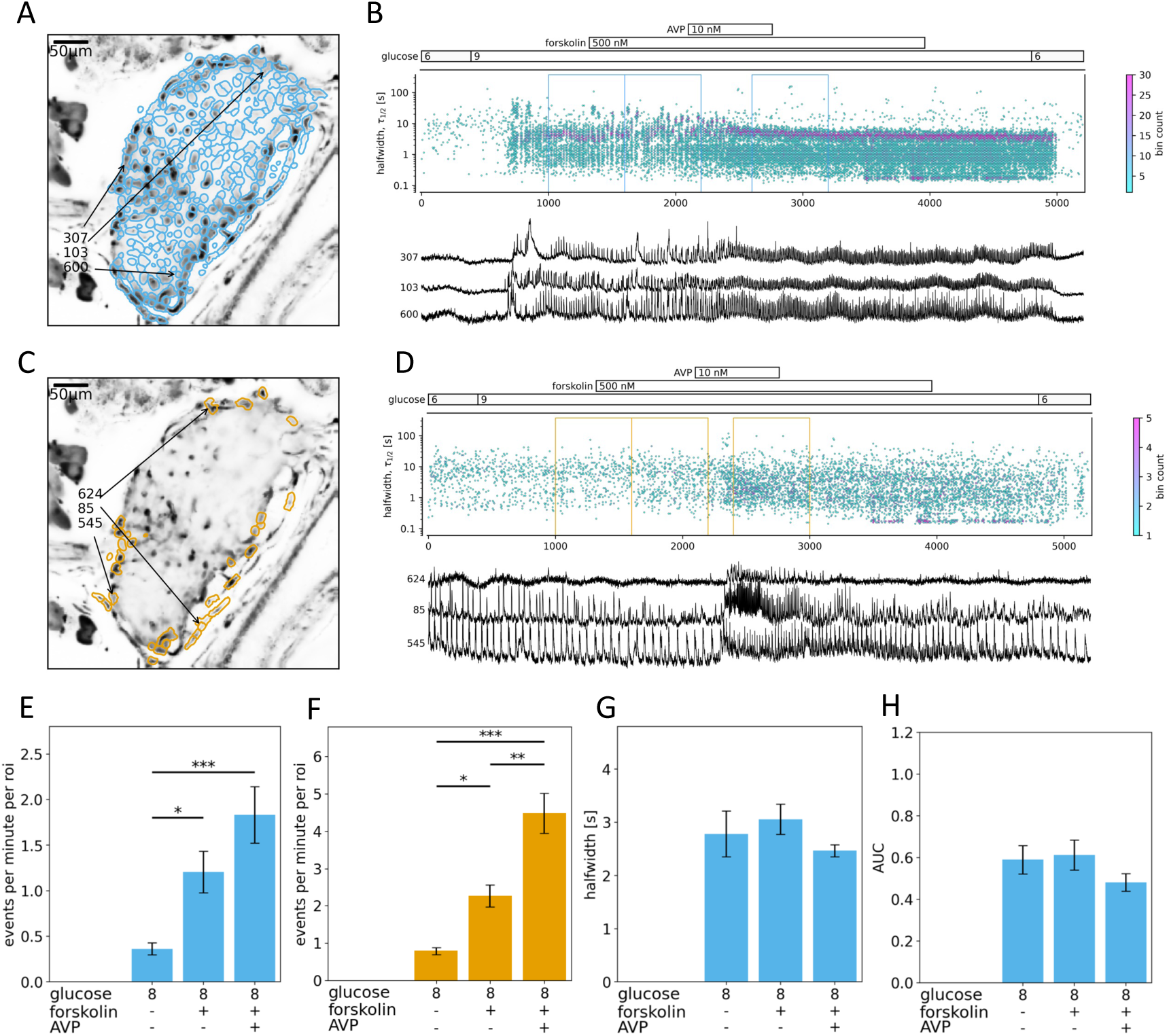
The effect of AVP in the physiological stimulatory glucose range on β (A, B, E, G, H) and α (C, D, F) cells. (A, C) Indicated are the ROIs whose filtered traces correlate best with the average trace for the whole islet. (B, D) (top panel) Time distribution of all measured eventś halfwidths at their peak times. Stimulation protocol is indicated above. The color bar indicates the bin count in each time/halfwidth point. The dashed-line rectangles indicate the 3 time periods where events per minute per ROI, halfwidth and AUC parameters were sampled for panels E-H. (bottom panel) Representative traces from ROIs indicated in A and C. (E, F) Forskolin significantly increased the number of Ca^2+^ oscillations per minute per ROI in β cells that were further stimulated by addition of AVP. A similar effect was observed also in α cells. (G, H) A non-significant but visible decrease in the events’ halfwidth and AUC values due to forskolin and AVP administration in β cells were noted. The parameters were distributed normally, and we used one-way ANOVA and Tukey post-hoc test. The data are presented as mean +/− SD (*p ˂ 0.05, **p ˂ 0.01, **p ˂ 0.001).

Using low levels of forskolin in the same islet increased the activity of α cells despite the presence of high glucose concentration and glucose-dependent Ca^2+^ oscillations in β cells, both of which should inhibit the activity of α cells (Fig. 3DF). This increased activity was further enhanced by adding 10 nM AVP (Fig. 3DF). Again, the intense stimulation of α cells had no detectable effect on AVP modulation of β cell activity. The control experiment without AVP application did not produce significant changes in oscillation frequency in a comparable recording time (Suppl. Fig. 3). In this study we did not observe any sex differences in AVP responses.

### The effect of physiological epinephrine concentration on AVP modulation of α and β cells

To further test whether AVP-dependent modulation of islet cell activity could be explained by a cAMP-like stimulatory mechanism, we used physiological epinephrine concentrations. We have previously shown that epinephrine suppresses Ca²⁺ activity in β cells at physiological glucose concentrations through a G_i_-protein-coupled pathway, while simultaneously stimulating α-cell activity through G_s_-protein-coupled signaling (51).

In the stimulatory 9 mM glucose, the administration of epinephrine rapidly reduced the frequency of β cell oscillations to a non-stimulatory activity level (Fig. 4). Subsequent perfusion with progressively increasing AVP concentration did not restore β cell activity beyond this level, indicating that AVP does not counteract epinephrine-induced inhibition in a manner consistent with activation of G_s_-protein-coupled V_2_ receptors (Fig. 4E).

**Figure 4.**
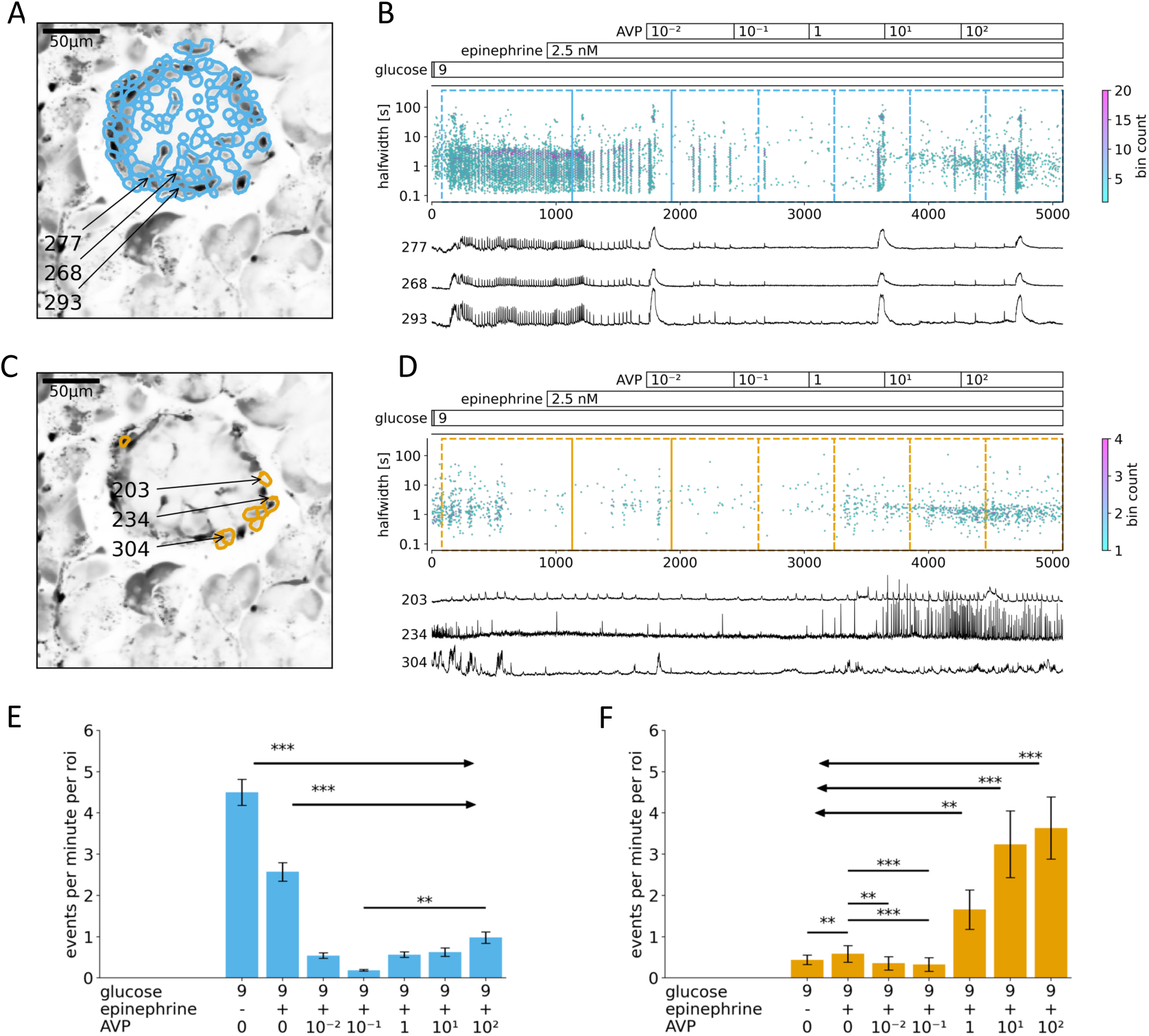
The effect of physiological epinephrine concentration β (A, B, E) and α (C, D, F) cells. (A, C) Indicated are the ROIs whose filtered traces correlate best with the average trace for the whole islet. (B, D) (top panel) Time distribution of all measured eventś halfwidths at their peak times. Stimulation protocol is indicated above. The color bar indicates the bin count in each time/halfwidth point. The dashed-line rectangles indicate the 7 time periods where events per minute parameter was sampled for panels E and F. (bottom panel) Representative traces from ROIs indicated in A and C. (E, F) The analysis of events per minute per ROIs shows that inhibition by epinephrine of glucose-dependent activation is reversed with increasing vasopressin concentration (0.01, 0.1, 1, 10 and 100 nM) in α cells but not in β cells. The parameters were distributed normally, and we used one-way ANOVA and Tukey post-hoc test. The data are presented as mean +/− SD (*p ˂ 0.05, **p ˂ 0.01, **p ˂ 0.001).

In contrast, α cells were activated by AVP and responded further to AVP in a frequency-coded epinephrine concentration-dependent manner (Fig. 4DF). This supports the view that AVP acts as a potent modulator of α cell activity under physiological adrenergic conditions. Importantly, these experiments were performed without forskolin, demonstrating that forskolin is not required for AVP-dependent activation of α cells or for detectable modulation of β cell activity. Rather, forskolin was used in subsequent experiments to provide a common cAMP-permissive background in which AVP effects on α and β cell activity could be compared under more standardized conditions.

### AVP enhances the activity of α and β cell within its physiological osmoregulatory range

To further investigate the sources of variability in the observed AVP action in the presence of forskolin and 8 mM glucose, we performed a series of experiments in which slices were exposed to an AVP concentration ramp. AVP administration reproducibly induced concentration-dependent activation of both α and β cells (Fig. 5). The most prominent stimulatory effect of AVP on both cell types was observed in its physiological osmoregulatory range between 10 and 100 pM (Fig. 5BFG).

**Figure 5.**
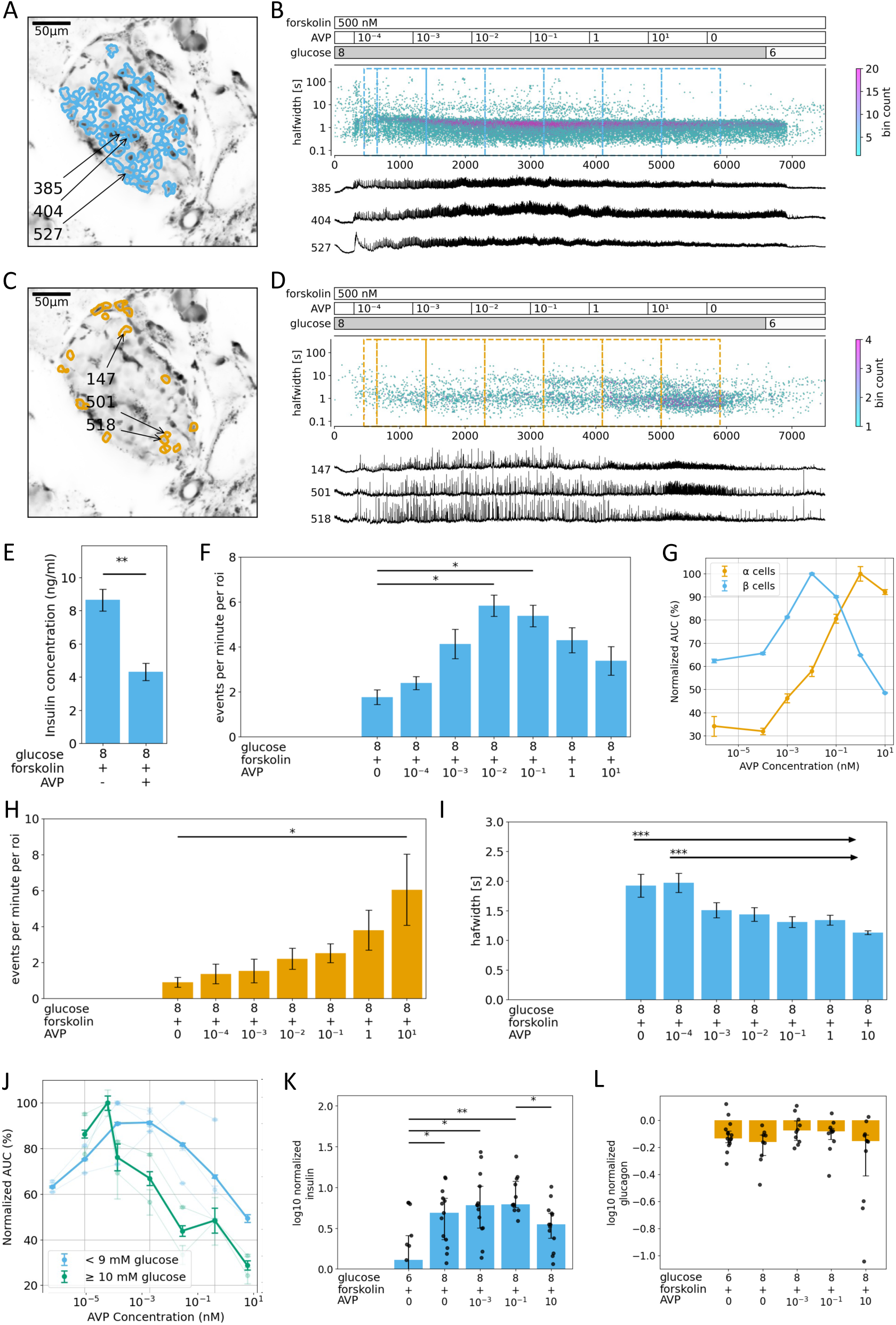
AVP modulates the activity of β (A, B, E-G, I, J, K) and α (C, D, G, H, L) cells in a concentration-dependent manner. (A, C) Indicated are the ROIs whose filtered traces correlate best with the average trace for the whole islet. (B, D) (top panel) Time distribution of all measured eventś halfwidths at their peak times. Stimulation protocol is indicated above. The color bar indicates the bin count in each time/halfwidth point. The dashed-line rectangles indicate the 7 time periods where events per minute per ROI and halfwidth parameters were sampled for panels F, H and J. (bottom panel) Representative traces from ROIs indicated in A and C. (E) Supraphysiological concentrations of AVP (above 10 nM) reduced insulin release in static incubation conditions from β cells below that measured at 8 mM glucose and forskolin alone. (F, H) Events per minute per ROI in β cells was gradually increased from 0.0001 nM to the physiological osmoregulatory range of AVP, between 0.01 and 0.1 nM, and then gradually decreased with further increases in AVP concentration. In the same islets the number of events per minute increased gradually in α cells also at higher AVP. (G) Pooling several islets revealed that bell-shaped distribution of the AUC could be found in both β and α cells, although the latter being shifted towards higher AVP concentration. (I) The halfwidth of events in β cells progressively declined with increased AVP concentration. (J) The AVP-dependent activation/inactivation curve for β cells was left-shifted at supraphysiological glucose concentration. All the parameters were distributed normally, and we used one-way ANOVA and Tukey post-hoc test. The data are presented as mean +/− SD (*p ˂ 0.05, **p ˂ 0.01, **p ˂ 0.001). (K, L) Concentration-dependent effects of AVP (in nM) on hormone secretion at 8 mM glucose and forskolin. (K) Insulin secretion exhibited a bell-shaped response, with maximal stimulation at low AVP concentrations and reduced secretion at 10 nM AVP. (L) Glucagon secretion was not significantly affected. Bars indicate median ± IQR; dots denote individual experiments. *P < 0.05, **P < 0.01.

In β cells, lower concentrations of AVP resulted in an increased frequency of Ca^2+^ oscillations, but this was reduced at higher AVP concentrations (Fig. 5G). In contrast, the halfwidth of oscillations decreased progressively and significantly with increasing AVP concentration (Fig. 5I). The overall effect of AVP produced a bell-shaped relationship for both the frequency parameter, measured as events per minute per region of interest (epmpr) (Fig. 5F), and normalized AUC values (Fig. 5G).

Such bell-shaped relationships are consistent with the known activation and inactivation properties of IP₃ receptors (37), which represent a major downstream component of V_1b_/G_q_/PLC-dependent Ca²⁺ signaling. IP_3_ receptors are activated with lower IP_3_ and Ca^2+^ levels, but become progressively inactivated at higher concentrations. Thus, unphysiologically high AVP concentrations may strongly stimulate of G_q_ protein-coupled signaling and thereby contribute to IP_3_ receptor inactivation, reducing the dynamic range of the AVP response and potentially resulting in an inhibitory effect, with AUC values falling below those of 8 mM glucose stimulation alone (Fig. 5G). This inhibitory effect was also reflected in the reduced insulin release in static incubation insulin release assay (Fig. 5E) as well as in dynamic insulin secretion (Fig. 5K). In the latter, insulin secretion showed a bell-shaped AVP concentration dependence, with increased release at low AVP concentrations and reduced release at 10 nM AVP.

A similar paradoxical inhibitory effect on β cells has previously been documented for muscarinic receptor signaling, which also involves the activation of Gq/PLC/IP₃ pathway (52). Additionally, at supraphysiological glucose concentration, the AVP-dependent activation/inactivation curve for β cells was shifted to the left (Fig. 5J), allowing for inhibitory modulation by AVP at lower concentrations and suggesting that glucose-dependent excitability state of β cells strongly shapes their sensitivity to AVP.

Notably, peak AVP activation in α cells occurred at an approximately one order of magnitude higher AVP concentration than in β cells (Fig. 5G), with limited evidence of inactivation at physiological AVP levels (Fig. 5GH), supporting previously published findings (13). Although AVP induced concentration-dependent changes in α cell Ca²⁺ activity, these effects were not reflected in dynamic glucagon secretion measurements. (Fig. 5L).

### V_1b_ receptor-selective agonists reproduce key AVP effects on α- and β-cell Ca²⁺ activity

One of the primary objectives of this study was to evaluate the exclusive role of V_1b_ receptors without confounding effects of the activation of V_1a_, V_2_ and oxytocin receptors in α and β cells using fresh pancreatic tissue slices. To accomplish this, we employed AVP, alongside a set of newly synthesized specific peptides (Fig. 6A): d[Cha^4^,Dab^8^]-VP, a specific V_1b_ receptor agonist; [(D-Leu)^2^,Ile^3^,Thr^4^]-VP, a mixed V_1b_ receptor agonist and V_1a_ and oxytocin receptor antagonist; and d[(CH_2_)_5_^1^,Tyr(Me)^2^,Dab^5^,Tyr^9^]-VP, specific V_1a_ receptor antagonist. Further, we used a commercially available compound tolvaptan to further substantiate the absence of V_2_ receptors in β cells – evidenced by our inability to reverse epinephrine inhibition with AVP (Fig. 4).

**Figure 6:**
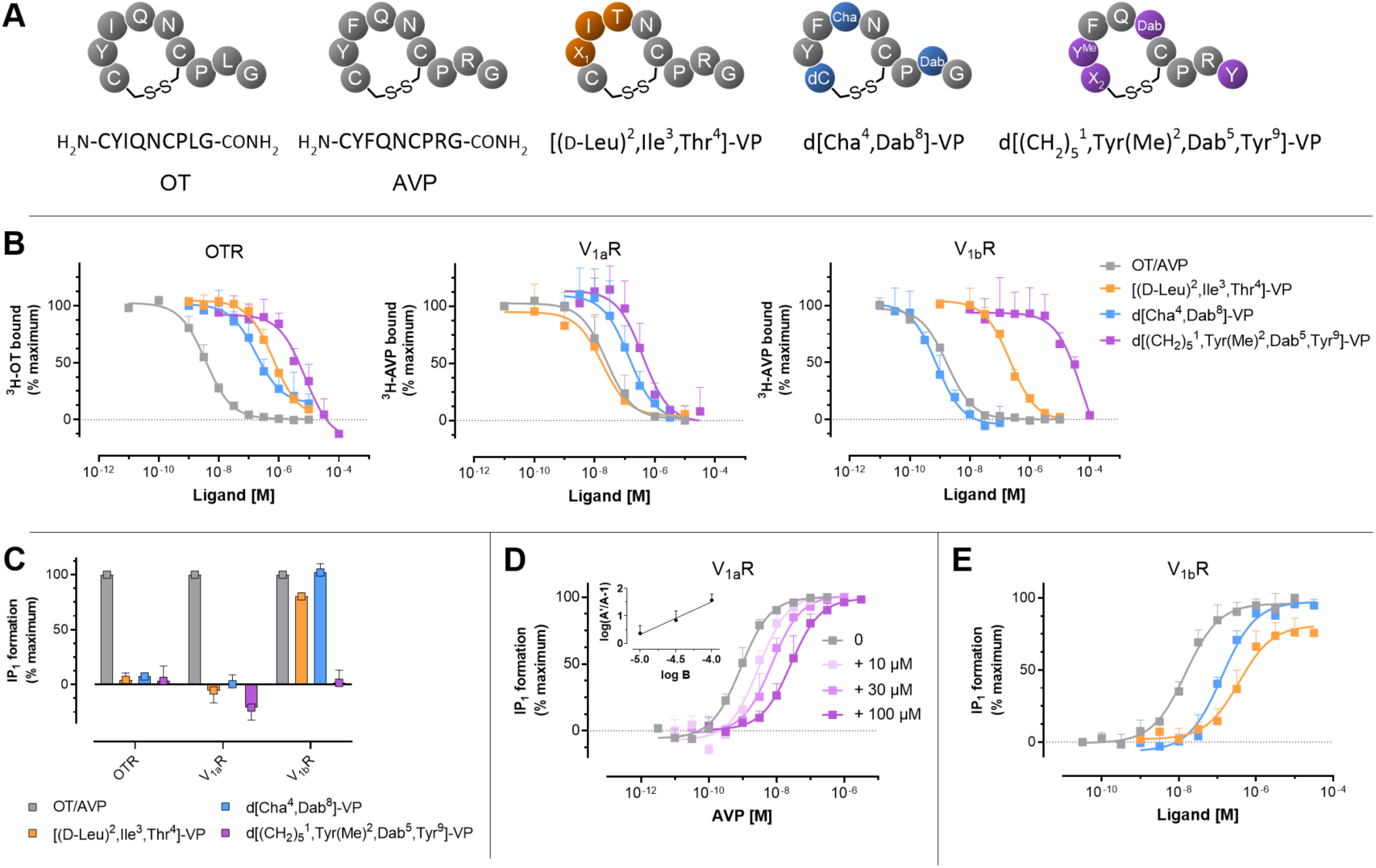
Schematic structures and pharmacology of peptide probes [(_D_-Leu)^2^,Ile^3^,Thr^4^]-VP, d[Cha^4^,Dab^8^]-VP, and d[(CH_2_)_5_^1^,Tyr(Me)^2^,Dab^5^,Tyr^9^]-VP. (A) Schematic structures and amino acid compositions of oxytocin (OT), vasopressin (AVP), and their derivatives. (B) Radioligand displacement assay of [(_D_-Leu)^2^,Ile^3^,Thr^4^]-VP, d[Cha^4^,Dab^8^]-VP, and d[(CH_2_)_5_^1^,Tyr(Me)^2^,Dab^5^,Tyr^9^]-VP. Concentration-dependent displacement of ^3^H-OT (0.65 nM) or ^3^H-AVP (0.21 nM for V_1a_R, 0.14 nM for V_1b_R) in HEK293 cell membranes expressing the human OTR, V_1a_R or V_1b_R by either test ligand or the respective control. Data is presented as specific binding by subtracting nonspecific from total binding normalized to the maximum binding of the radioligand in the absence of the peptides (n = 3). (C) Ligand induced formation of intracellular IP_1_ by 10 µM OT/AVP, [(_D_-Leu)^2^,Ile^3^,Thr^4^]-VP, d[Cha^4^,Dab^8^]-VP, and d[(CH_2_)_5_^1^,Tyr(Me)^2^,Dab^5^,Tyr^9^]-VP demonstrating agonistic properties of [(_D_-Leu)^2^,Ile^3^,Thr^4^]-VP and d[Cha^4^,Dab^8^]-VP at V_1b_R, whereas in the sense of an antagonism no increased IP_1_ accumulation was observed at OTR and V_1a_R (n = 2). Activation data was normalized to maximum response generated by the control (OT for OTR, AVP for V_1a_R and V_1b_R). All data points are shown as mean ± SD. (D) Antagonistic properties of d[(CH_2_)_5_^1^,Tyr(Me)^2^,Dab^5^,Tyr^9^]-VP at V_1a_R were assessed through concentration-dependent displacement of AVP according to the Schild method (n=3). (E) Concentration-response curves of AVP, [(_D_-Leu)^2^,Ile^3^,Thr^4^]-VP, and d[Cha^4^,Dab^8^]-VP were obtained for the V_1b_R in order to assess their potency (n = 3). Affinity constants (K_i_), potency (EC50) and maximum efficacy (E_max_) values for [(_D_-Leu)^2^,Ile^3^,Thr^4^]-VP, d[Cha^4^,Dab^8^]-VP, d[(CH_2_)_5_^1^,Tyr(Me)^2^,Dab^5^,Tyr^9^]-VP, and the respective controls are listed in Table 1. Abbreviations: Cha = cyclohexylalanine; Dab = 2,4-diaminobutyric acid; d = desamino; X_1_ = _D_-Leu; X_2_ = d(CH_2_)_5_ (β-mercapto-β,β-cyclopentamethylenepropionic acid); Y^Me^ and Tyr(Me)= O-methylated tyrosine.

The three peptide ligands were rationally designed based on available earlier structure-activity relationship data from our own previous work (47) and others (53–55), incorporating specific modifications to optimize receptor selectivity and activation. The functional properties of the three peptide ligands were assessed using a panel of cell-based *in vitro* assays expressing human oxytocin, V_1a_, and V_1b_ receptors. Receptor-subtype selectivity was assessed using radioligand binding studies (Fig. 6B) measuring the displacement of tritiated OT/AVP with increasing concentrations of peptides. This demonstrated that d[Cha^4^,Dab^8^]-VP had the highest affinity for V_1b_ receptor (200 pM) and a selectivity gain over oxytocin (∼270-fold) and V_1a_ receptor (∼115-fold) (Table 1). On the other hand, [(D-Leu)^2^,Ile^3^,Thr^4^]-VP had comparable affinity for all three receptors, ranging between 5 nM and 209 nM (Table 1), and lastly d[(CH_2_)_5_^1^,Tyr(Me)^2^,Dab^5^,Tyr^9^]-VP exhibited only affinity for V_1a_ receptor (66 nM), but did not bind to oxytocin (>2.3 µM) or V_1b_ receptors (>10 µM) (Table 1).

To determine their activation profile, intracellular signaling was evaluated using IP_1_ second messenger quantification in two steps: a first single-concentration screen allowed to distinguish agonist/antagonist properties of the peptide ligands (Fig. 6C) and a comprehensive concentration-response determination yielded reliable values for potency (EC_50_), efficacy (E_max_) and inhibitory potency (pA_2_) (Fig. 5DE). Peptide d[Cha^4^,Dab^8^]-VP is a moderate potent full agonist at V_1b_ receptor (EC_50_ = 130 nM; E_max_ = 104%) (Table 1), with no significant activation of V_1a_ or oxytocin receptors (Fig. 5C). Its potency was slightly reduced as compared to endogenous AVP (15 nM). Peptide [(D-Leu)^2^,Ile^3^,Thr^4^]-VP functioned as a moderate/weak partial agonist at V_1b_ receptor (EC_50_ = 361 nM; E_max_ = 79%) (Table 1), while displaying antagonist properties at oxytocin and V_1a_ receptors (Fig. 6C). Peptide d[(CH_2_)_5_^1^,Tyr(Me)^2^,Dab^5^,Tyr^9^]-VP demonstrated selective antagonism for V_1a_ receptor, effectively inhibiting endogenous ligand-induced signaling in a competitive manner (as exemplified by Schild regression analysis, Fig. 6D). Its inhibitory potency (pA2) was determined to 5.27 (Table 1), corresponding to 5.37 µM. These findings demonstrate that the newly designed peptide ligands provide valuable tools for probing the role of V_1b_ receptors in in α and β cells.

Applied in pancreatic tissue slices, [(D-Leu)^2^,Ile^3^,Thr^4^]-VP produced effects similar to AVP, characterized by an increase in the frequency and a decrease in the halfwidth of the Ca^2+^ oscillations (Fig. 7). However, paired before-after analysis revealed heterogeneous β cell responses, with some islets showing no change or even a reduction in Ca²⁺ oscillation frequency after ligand application. Consequently, the pooled AUC data did not show a significant difference (Fig. 7E). In contrast, α cells were activated in a frequency-coded manner (Fig. 7DF).

**Figure 7.**
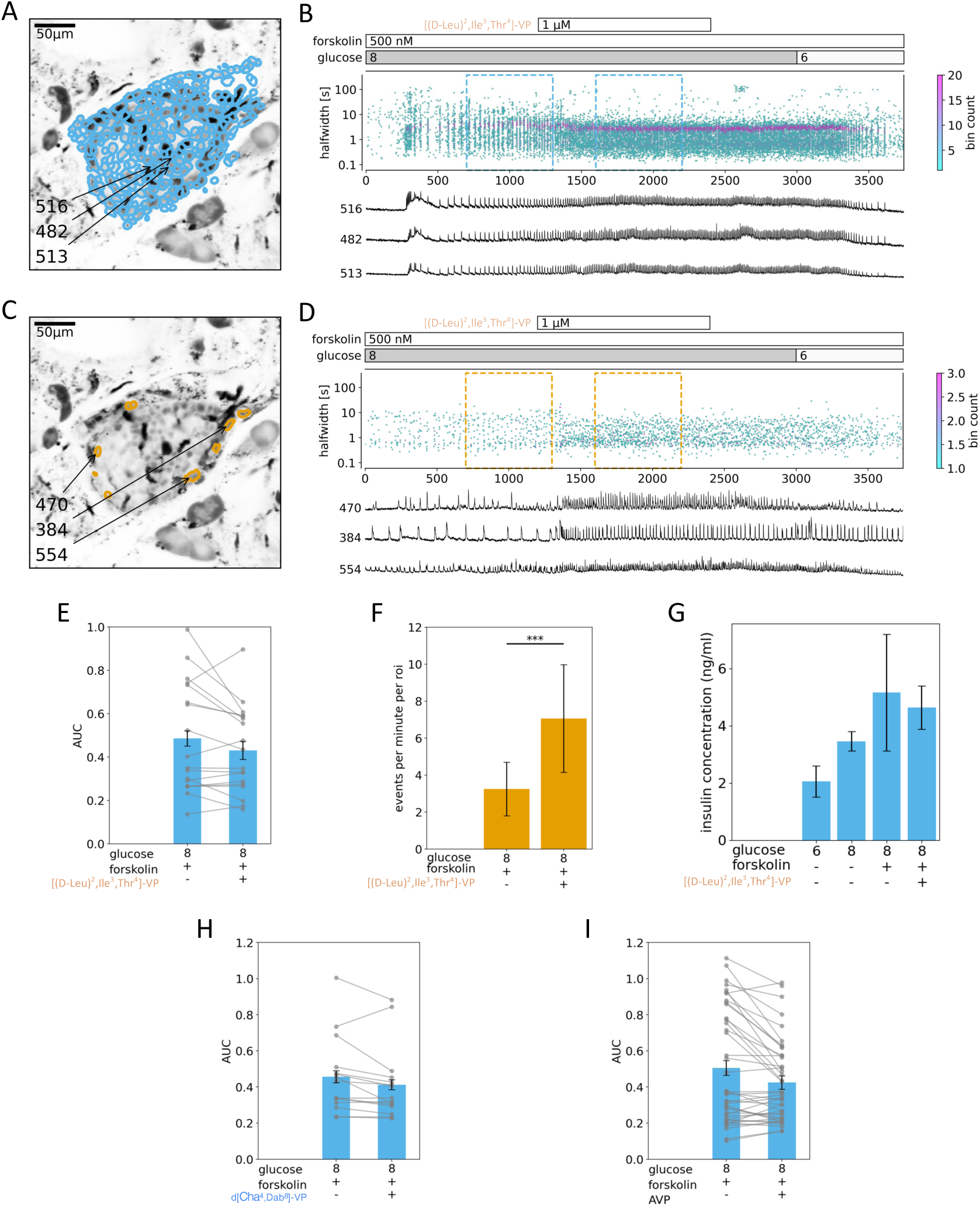
A specific V_1b_ receptor agonist and V_1a_ and oxytocin receptor antagonist, has a similar effect on β (A, B, E) and α (C, D, F) cells as AVP. (A, C) Indicated are the ROIs whose filtered traces correlate best with the average trace for the whole islet. (B, D) (top panel) Time distribution of all measured eventś halfwidths at their peak times. Stimulation protocol is indicated above. The color bar indicates the bin count in each time/halfwidth point. The dashed-line rectangles indicate the 2 time periods where events per minute per ROI and AUC parameters were sampled for panels E and F. (bottom panel) Representative traces from ROIs indicated in A and C. (E) There were no significant differences in the pooled AUC data for β cells treated with the [(D-Leu)^2^,Ile^3^,Thr^4^]-VP and forskolin or with forskolin only. Direct comparison of the AUC values before and after the ligand application shows a large heterogeneity of responses. (F) A significant increase in the events per minute per ROIs due to [(D-Leu)^2^,Ile^3^,Thr^4^]-VP administration in α cells was noted. (G) There was no significant effect on insulin release from slices after [(D-Leu)^2^,Ile^3^,Thr^4^]-VP exposure in comparison to stimulatory forskolin only. (H, I) The d[Cha^4^, Dab^8^]-VP (a specific V_1b_ receptor agonist) and AVP produced similarly heterogeneous responses in β cells with non-significant changes in the AUC compared to forskolin only. All the parameters were distributed normally, and we used one-way ANOVA and Tukey post-hoc test. The data are presented as mean +/− SD (*p ˂ 0.05, **p ˂ 0.01, **p ˂ 0.001).

Insulin release from islets within pancreas slices did not show statistically significant differences following treatment with [(D-Leu)^2^,Ile^3^,Thr^4^]-VP in 8 mM glucose and substimulatory concentration of forskolin (Fig. 7G). Given that [(D-Leu)^2^,Ile^3^,Thr^4^]-VP also antagonized V_1a_ and oxytocin receptors, it is plausible that the stimulatory effect through V_1b_ receptors was attenuated. Analysis of the effects of d[Cha^4^,Dab^8^]-VP or AVP under similar experimental conditions, revealed that both produced comparably heterogenous responses. Paired analysis indicated a non-significant change in AUC with both V_1b_ receptor-selective agonists (Fig. 7EH) as well with AVP (Fig. 7I).

Subsequently, we tested the involvement of V_1a_ receptors using d[(CH_2_)_5_^1^,Tyr(Me)^2^,Dab^5^,Tyr^9^]-VP, a V_1a_ receptor antagonist. This antagonist did not inhibit AVP-stimulated Ca^2+^ activity in either α and β cells (Fig. 8A-F). To validate its V_1a_ antagonistic action in the same tissue preparation, we assessed Ca^2+^ oscillations in smooth muscle cells of blood vessels adjacent to the islets (Fig. 8G). AVP increased the frequency of Ca^2+^ oscillations in vascular smooth muscle cells, and this activity was specifically inhibited by d[(CH_2_)_5_^1^,Tyr(Me)^2^,Dab^5^,Tyr^9^]-VP and restored when antagonist washout, consistent with AVP action through V_1a_ receptor in these cells. In contrast, application of V_2_ receptor antagonist tolvaptan did not have any significant effect on α cells, β cells, or any other cell types in the slice, and did not reverse any of the AVP effects (Suppl. Fig. 2).

**Figure 8.**
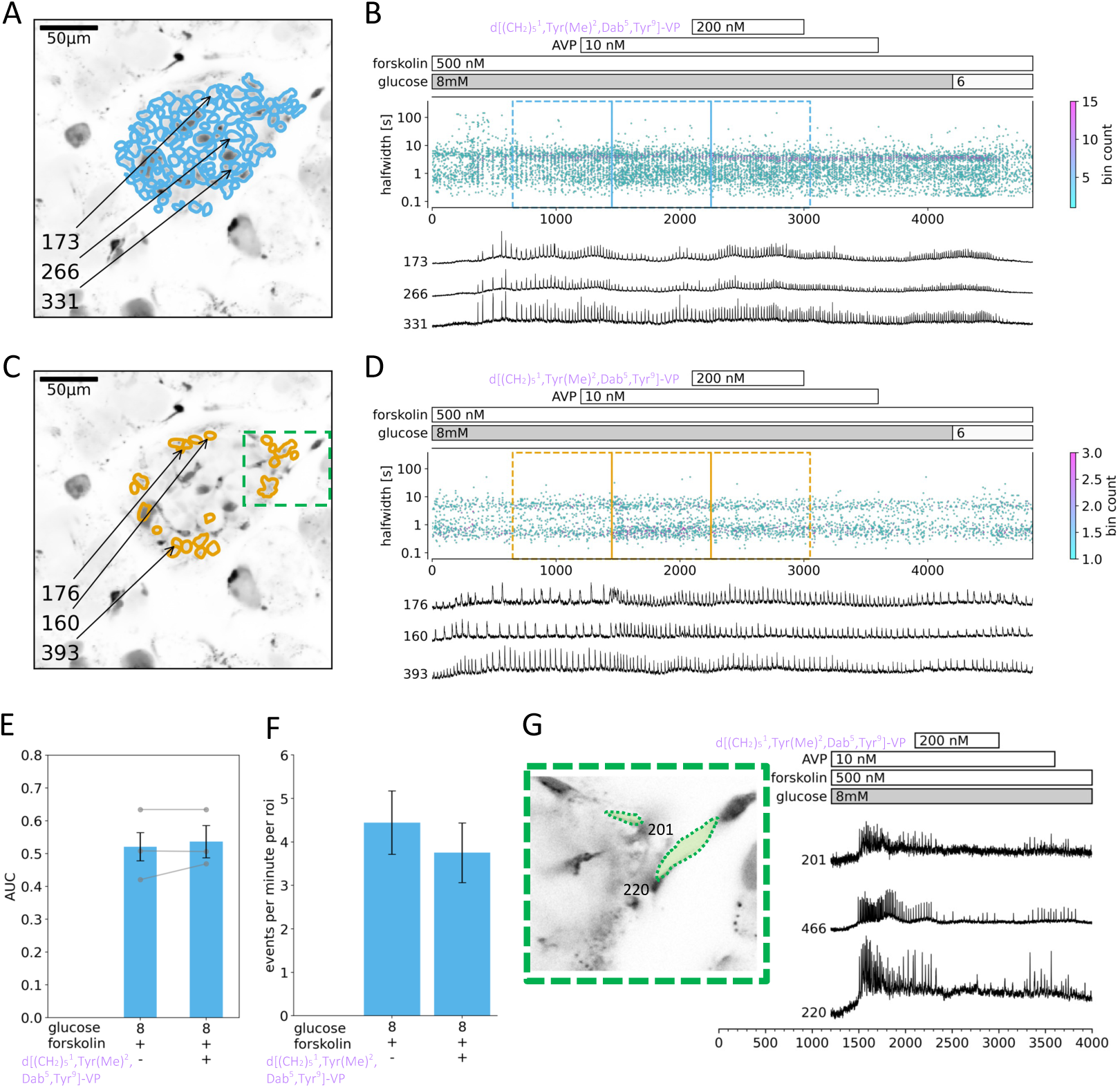
A specific V_1a_ receptor antagonist has no significant effect on β (A, B, E, F) and α (C, D) cells, but inhibits activity in smooth muscle cells (G). (A, C) Indicated are the ROIs whose filtered traces correlate best with the average trace for the whole islet. Inset labelled with green dashed line is expanded in panel G. (B, D) (top panel) Time distribution of all measured eventś halfwidths at their peak times. Stimulation protocol is indicated above. The color bar indicates the bin count in each time/halfwidth point. The dashed-line rectangles indicate the 3 time periods where events per minute per ROI and AUC parameters were sampled for panels E and F. (bottom panel) Representative traces from ROIs indicated in A and C. (E) There were no significant differences in the pooled AUC data for β cells treated with the d[(CH_2_)_5_^1^, Tyr(Me)^2^, Dab^5^, Tyr^9^]-VP and forskolin or with forskolin only. Direct comparison of the AUC values before and after the antagonist application shows no difference in responses. (F) No significant differences in the events per minute per ROIs due to d[(CH_2_)_5_^1^, Tyr(Me)^2^, Dab^5^, Tyr^9^]-VP administration in α cells were noted. (G) (left) Inset from C, showing location of two smooth muscle cells lining the blood vessel adjacent to the islet (right) Representative traces from ROIs on smooth muscle cells exposed to stimulatory glucose, forskolin, AVP, and d[(CH_2_)_5_^1^, Tyr(Me)^2^, Dab^5^, Tyr^9^]-VP. AVP administration increased the frequency of Ca^2+^ oscillations in smooth muscle cells, but V_1a_ receptor antagonist fully, but reversibly inhibited this activity. All the parameters were distributed normally, and we used one-way ANOVA and Tukey post-hoc test. The data are presented as mean +/− SD.

### State-dependent pharmacological modulation of β cell activity by AVP and V_1b_ receptor-selective agonists

Fig. 9 summarizes the analysis of the pharmacological modulation of AVP receptor-dependent responses in β cells within pancreatic tissue slices. During the AVP concentration ramp, both the amplitude and direction of changes in the rate of Ca²⁺ oscillations varied substantially across islets (Fig. 9A). The initial activation at lower pM AVP concentrations was followed by a relatively stable response near the peak of a bell-shaped concentration-dependence curve, which then transitioned towards inhibition at concentrations approaching or exceeding nM range.

**Figure 9.**
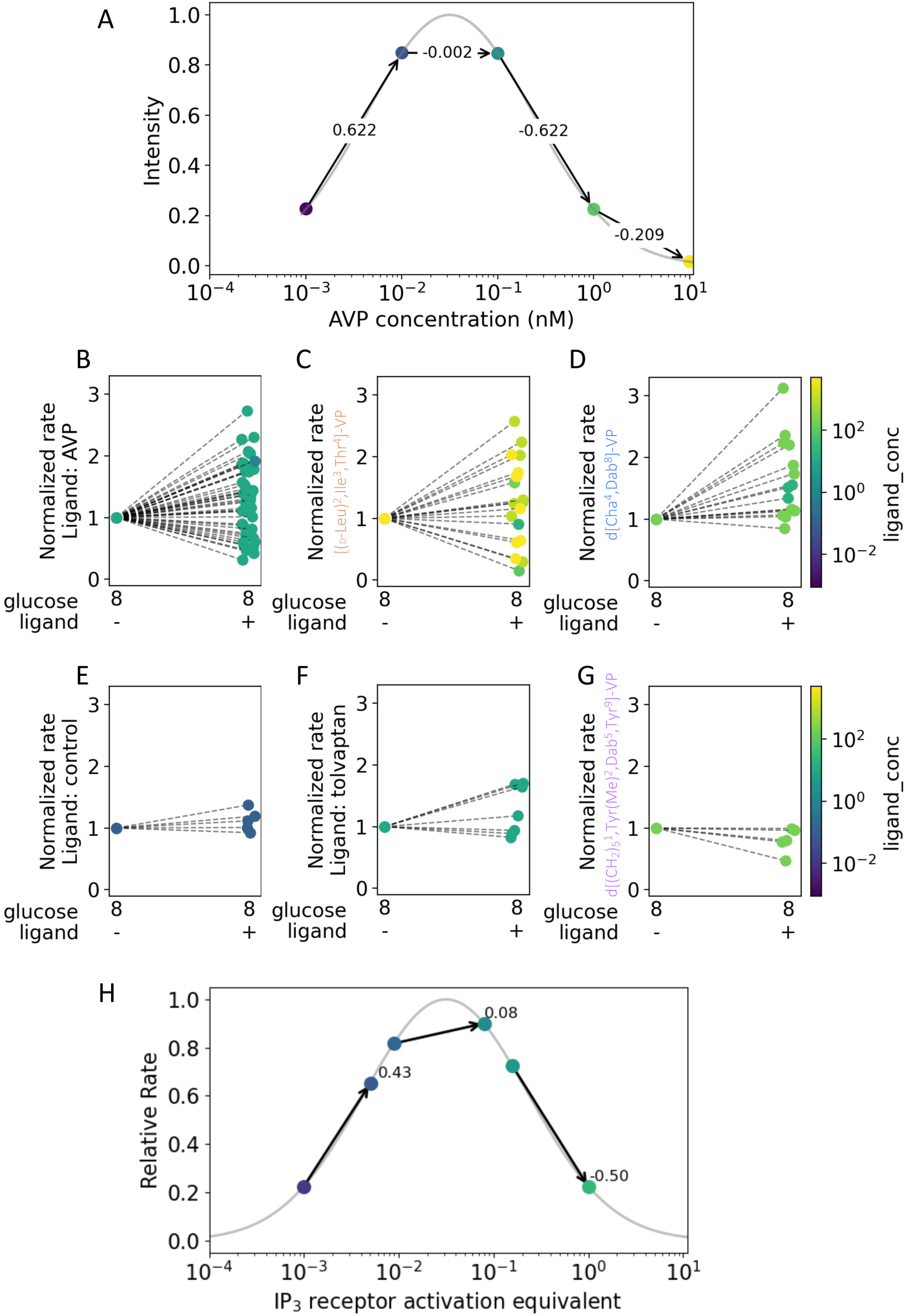
Overview of the analysis of pharmacological modulation of V_1b_ receptors in β cells in pancreatic tissue slices. (A) The relative rates of Ca^2+^ oscillations obtained during the AVP concentration ramp experiment. The bell-shaped dependence curve rises at the lower pM range of AVP, followed by a relatively stable response around the peak of the curve, and then falls at higher concentrations (nM range) of AVP. (B, C, D) Normalized rates to the basal stimulatory conditions (8 mM glucose) in the context of single concentration static incubation with AVP, [(D-Leu)^2^, Ile^3^, Thr^4^]-VP and d[Cha^4^,Dab^8^]-VP are widely scattered. (E, F, G) Normalized rates to the basal stimulatory conditions (8 mM glucose) in the context of single concentration static incubation without AVP receptor agonists (control), with tolvaptan (V_2_ antagonist) and d[(CH_2_)_5_^1^,Tyr(Me)^2^,Dab^5^,Tyr^9^]-VP (V_1a_ antagonist) are only slightly scattered. (H) Different islets during their plateau phase stimulated by 8 mM glucose and forskolin can achieve different levels of IP_3_ receptor activation equivalent, contributing to the heterogeneity of further IP_3_R stimulation with AVP or specific V_1b_ agonists.

The distribution of the rate of changes at different AVP concentrations resembled that observed during static incubation experiment with AVP or V_1b_ receptor-selective agonists (Fig. 9B), showing a range of responses: activation (46%), no effect (27%), and inhibition (17%) after applying AVP, [(D-Leu)^2^,Ile^3^,Thr^4^]-VP (Fig. 8C) or d[Cha^4^,Dab^8^]-VP (Fig. 8D). In contrast, control without agonists (Fig 8E) or the use of antagonists for non-expressing AVP receptors following AVP stimulation, did not significantly affect the relative rate change (Fig. 8FG). One interpretation of the observed variability with V_1b_ receptor activation is that different islets, during their plateau phase stimulated by 8 mM glucose and forskolin, may achieve varying levels of IP_3_ receptor activation (IP_3_ receptor activation equivalent), before being further influenced by AVP or specific V_1b_ receptor agonists (Fig. 8H). Overall, the effect of V_1b_ receptor stimulation depends on the current functional status of an islet. Resting β cells permit activation and increase insulin secretion, whereas in highly activated β cells, high levels of AVP and V_1b_ receptor agonists tend to promote inactivation and interfere with insulin release (Fig. 5K).

## Discussion

The use of freshly prepared pancreatic tissue slices was important for resolving the context-dependent role that AVP receptor signaling plays in the physiology of pancreatic β and α cells. This preparation represents acutely isolated, viable pancreatic tissue from healthy mice that can be imaged in real time using high-resolution confocal microscopy shortly after extraction. Tissue manipulation during slicing and vital Ca²⁺ indicator loading is kept to a minimum. Pancreatic slices therefore provide a valuable ex vivo preparation that preserves many properties of intact cells, including local tissue architecture and intercellular communication between homotypic and heterotypic cell types. These interactions are disrupted in research models involving isolated islets, dispersed islet cells (56) and cell lines, and the slice preparation also avoids the enzymatic digestion step used for islet isolation (38, 57). At the same time, pancreatic slices remain an ex vivo model with limitations, including the absence of intact vascular perfusion and innervation.

Existing evidence regarding the role of AVP for the activity of pancreatic β and α cells remains controversial. Some studies suggest that AVP plays a significant role in insulin and glucagon release, whereas others report minimal or selective effects (7, 13). There has generally been stronger functional evidence for involvement of V_1b_ receptors than direct evidence for their mRNA or protein abundance in β cells. Previous single-cell transcriptomic analysis (58) or protein expression analysis (8) reported absent or very low AVP receptor expression in β cells, whereas overexpression of V_1b_ receptors in β cells improved the function of transplanted pseudoislets in vitro (59). Our new RNAscope analysis adds spatial information to this discussion (Fig. 1). It confirms that V_1b_ receptor transcripts are present in glucagon-positive α cells, but also shows that V_1b_ receptor transcript signal in the islet is not restricted to α cells. In our dataset, 67.9 ± 7.5% of α cells were V_1b_ receptor-positive, whereas only approximately one third of the total V_1b_ receptor-positive signal/cells colocalized with glucagon-positive α cells. Thus, the data support α cells as an important V_1b_ receptor-positive endocrine population, but argue against a strictly α-cell-restricted model of V_1b_ receptor expression in intact pancreatic tissue (28). Future approaches combining spatial expression analysis, cell-specific genetic models, and physiological recordings in intact tissue will be needed to determine how AVP responses arise from direct receptor expression, α-to-β-cell interactions (60), and the dynamic state of the β-cell collective.

Our research aimed to reassess the role of AVP in the function of both α and β cells in fresh pancreas tissue slices, a preparation with in vivo-like sensitivity to physiological and pharmacological stimuli while enabling high spatial and temporal resolution measurements (38, 49, 51, 61). We focused on signaling through G_q_-coupled AVP receptor, particularly V_1b_ receptors, and downstream Ca^2+^ signaling, involving IP_3_ receptors. The bell-shaped response curve observed with AVP receptor agonists, where lower concentrations stimulate and higher concentrations inhibit cell activation and insulin release, is consistent with the biphasic activation and inactivation properties of the IP_3_ receptors (37, 62, 63). The single-concentration agonist experiments with 15-minute incubations allowed us to reduce the possibility that the observed responses during the prolonged AVP ramp were explained solely by time-dependent receptor internalization or desensitization. The bell-shaped relationship has important implications for understanding how β and α cells respond to fluctuating AVP levels of AVP under varying plasma glucose levels, plasma osmolality and hormonal context. Importantly, it underscores the necessity of controlling AVP to avoid dysfunctional endocrine responses, or improve it after transplantation (59). This interpretation is also consistent with the emerging clinical view that AVP, as a hydration-sensitive hormone, links osmoregulatory state to metabolic outcomes, including glucose homeostasis, insulin sensitivity, lipid metabolism, and energy balance(64).

Hormones acting through IP_3_ as a Ca^2+^-releasing messenger can induce intracellular Ca^2+^ oscillations that are shaped by Ca^2+^ feedback on the IP_3_ receptor (65). We combined high-resolution measurements of cytosolic Ca^2+^ oscillations and a high-throughput analysis of newly synthesized V_1b_ receptor-selective ligands, including ligands that simultaneously antagonized V_1a_ and oxytocin receptors.Our findings support a model in which AVP modulates the activity of β and α cells in a non-linear, state-dependent manner. The observed effects across the physiological AVP concentration range, with IP_3_ receptor-dependent signaling as a functional downstream readout, followed a bell-shaped activation/inactivation profile. This non-linear dependence may help explain why previous studies reached conflicting conclusions, particularly when experiments were performed under different glucose, cAMP, or hormonal conditions, or were interpreted without considering the activation/inactivation profile previously observed in perfused rat pancreas (18) and insulinoma cells (29).

By measuring the AVP-dependent activity of both cell types simultaneously, our results do not support a simple model in which activation of V_1b_ receptors in α cells is sufficient to indirectly activate β cells in a coordinated paracrine fashion. Rather, β and α cell responses were temporally and functionally distinct, suggesting that β cell modulation cannot be explained solely as a secondary consequence of α cell activation. Nevertheless, paracrine interactions within the intact islet cannot be excluded and may contribute to the overall response under specific conditions, for example possible direct AVP activation of δ cells that would in turn inhibit insulin release from β cell. However, a qualitatively similar bell-shaped dependence has also been described for β cell responses to acetylcholine stimulation, although interpreted differently (66). Together, these findings support the interpretation that AVP acts less as a simple on/off stimulus and more as a context-dependent perturbation of the islet Ca²⁺ signaling network.

### Glucose-Dependent Modulation by AVP

Our results reproduced the glucose-dependent effects of AVP on both α and β cells, corroborating previous studies. One important differences from most previous work is that AVP reached its peak activity within its physiological osmoregulatory range of 10-100 pM (1), rather than in the nM range. We showed that AVP-dependent augmentation of β cell activity required stimulatory glucose concentrations. In contrast, AVP was able to activate α cells even at glucose-concentrations that are typically inhibitory for these cells, particularly when cAMP levels were elevated either directly by forskolin-mediated stimulation of adenylyl cyclase or physiologically by epinephrine acting through G_s_-protein-coupled β-adrenergic receptors. Furthermore, simultaneous α and β cell activity, enabled by forskolin, revealed no major α-β cell-cell interaction at the time scales analyzed in this study. β cell were more sensitive to AVP, with peak activity between 10 and 100 pM, whereas α cell activity peaked at approximately 1 nM AVP. It remains to be investigated whether this apparent lack of α-β cell communication reflects the fact that AVP primarily drives Ca^2+^ release through IP_3_ receptors, rather than through plasma membrane-related Ca^2+^ fluxes.

Even moderately supraphysiological glucose concentrations shifted the AVP concentration dependence of β cells to the left, such that physiologically relevant concentration of AVP were more likely to exert inhibitory effect on the activity of these cells. This suggests that, during hyperglycemia and increased plasma osmolality the rise in AVP concentration with concomitant elevation of cytosolic cAMP concentration through GLP-1 or other G_s_-protein-coupled receptor stimulation, could diminish the capacity of β cells to respond adequately, or even suppress glucose-dependent insulin release (64). Similarly, a glucose-dependent modulation of AVP sensitivity could affect the activation capacity of α cells in hypoglycemic conditions and contribute to the inability of α cells to counteract hypoglycemia and contribute to impaired counterregulatory responses. The rightward shift in AVP dependence in α cells compared with β cells may reflect a generally lower activation state under glucose concentrations that are suboptimal for α cell activity, and this sensitivity may shift under more favorable glucose conditions. Further experiments will be required to clarify this issue.

It is important to note that although AVP can potentiate α cell activity, this potentiation has been reported to be insufficient in T1D patients to support an adequate counterregulatory response during hypoglycemia, a context associated with high systemic AVP concentrations (13). Overactivation of α cells by AVP in the presence of high cAMP levels, for example during concomitant stress-related adrenergic stimulation, could potentially desensitize or impair the ability of α cells to respond appropriately to dangerously low glucose concentrations. Thus, under certain pathological or extreme physiological conditions, AVP together with elevated cAMP may disrupt normal glucagon release and compromise the capacity to counteract hypoglycemia.

### Permissive Role of AVP in cAMP-Dependent Signaling

A critical aspect of AVP’s role in β and α cells is its dependence on the prevailing cAMP signaling environment. Our study provides evidence that AVP effects were strongly shaped by intracellular cAMP environment, with forskolin enabling the simultaneous assessment of AVP responses in both cell types, despite their differential sensitivity to glucose. This suggests that AVP does not act primarily as a universal initiator of β and α cell activity, but rather enhances or reshapes the response of these cells to other stimuli when cAMP levels are elevated. The permissive role of AVP, particularly in the context of cAMP-dependent signaling, aligns with the broader understanding of its function in other endocrine systems, such as CRH- or histamine-dependent potentiation of ACTH release in the pituitary (32, 34).

The observation that epinephrine, which reduces cAMP levels and suppresses Ca^2+^ activity in β cells, did not allow AVP to restore β cell activity beyond the resting level further supports the interpretation that AVP action is closely linked to the cAMP signaling axis. This cell-type-specific interaction may help explain the variability in AVP effects observed across different studies and species, where differences in glucose concentration, adrenergic tone, incretin signaling, or experimental cAMP background could strongly modulate the apparent AVP response. Consistent with this permissive framework, antagonism of GLP-1 receptors prevented AVP-dependent action in β cells (58).

### V_1b_ Receptor-Specific Modulation

The heterogeneous responses observed across different islets, where some showed increased activity while others showed no detectable change or inhibition, could intuitively be attributed to variability in V_1b_ receptor expression or signaling capacity among β cells. Such an explanation would be consistent with differences in receptor density, coupling efficiency to G_q_ proteins, or downstream signaling components such as PLC or IP₃ receptors. However, our data suggest that receptor-level variability alone is unlikely to fully explain the observed response spectrum, and that the current functional state of the islet collective must also be considered. The islet behaves as a non-linear dynamic system in which the same molecular perturbation can produce different functional outcomes depending on the current state of the β cell collective. In such systems, cells or cell populations do not occupy a single deterministic activity state, but rather move within a landscape of possible states, with perturbations shifting the probability distribution of transitions between them (67). This concept is well established in dynamical systems approaches to biological cell-state transitions, where attractor landscapes, noise, and signaling inputs determine the probability of moving between alternative functional states rather than enforcing a single fixed output.

In this framework, AVP and V_1b_ receptor-selective agonists may reshape the probability landscape of β cell activity. Depending on the initial metabolic, electrical, and Ca²⁺-handling state of the islet, the same stimulus may increase oscillation frequency, produce little detectable effect, or shift the system toward reduced activity or functional inactivation. This interpretation is also consistent with studies of pancreatic islet dynamics showing that β cell Ca²⁺ activity emerges from coupled electrical, metabolic, and network interactions rather than from the properties of individual cells alone (68). Thus, molecular variability in V_1b_ receptor expression or signaling capacity may contribute to the heterogeneous responses, but it is unlikely to determine them without considering the collective dynamic state of the islet.

The study specifically targeted the V_1b_ receptor to distinguish its role from other AVP receptor subtypes that have been previously described to contribute to islet cell responses, including V_1a_, V_2_ and oxytocin receptors. The use of V_1b_ receptor-selective agonists together with V_1a_ and oxytocin receptor antagonists allowed a more precise characterization of AVP modulation of β and α cell function. The findings support V_1b_ receptor-dependent signaling as a major mediator of AVP effects in pancreatic tissue slices, with no detectable contribution from V_1a_, V_2_ or oxytocin receptors under the conditions tested. In addition, previous functional studies using selective V_1b_ receptor agonists, such as d[Cha4]AVP, revealed nanomolar affinity for V_1b_ receptors across species and concentration-dependent activation of natural biological models expressing these receptors (69).

### Physiological Implications and Broader Context

Maintaining AVP levels within a physiological range is particularly challenging in diabetes mellitus. Hyperglycemia can increase plasma osmolality and promote water loss, whereas hypoglycemia can also directly stimulate AVP release (13). In these scenarios, AVP signaling becomes physiologically relevant not only as an osmoregulatory response, but also as a potential modulator of α and β cell activity(64). Elevated AVP may therefore contribute to endocrine dysregulation when the islet is already exposed to altered glucose, cAMP, and stress-related signaling conditions.

The complex interplay between AVP, glucose concentration, and the endocrine state of the islet also highlights a broader conceptual issue. AVP does not appear to impose a single deterministic response on β cells. Rather, as summarized in the graphical abstract, the same AVP stimulus may shift an islet between different activity states depending on its initial position within a non-linear functional landscape. At 8 mM glucose, a weakly active β cell collective may remain responsive and be further activated by AVP, whereas a highly active collective may be pushed toward a damped or trapped state. This state dependence provides a framework for understanding why AVP can stimulate, fail to affect, or inhibit β cell activity under apparently similar experimental conditions. In a diabetes, where AVP levels may be elevated because of chronic hyperosmolarity, dehydration or hypoglycemia, the probability of tipping the islet towards an inhibitory or desynchronized state may increase. This could impair insulin secretion and, depending on the glucose and cAMP context, may also disturb the glucagon response during hypoglycemia. Thus, AVP signaling should be considered not simply as an endocrine stimulus, but as a context-dependent perturbation that reshapes the probability distribution of accessible islet activity states.

Given these complexities, therapeutic strategies targeting AVP signaling in diabetes would require a nuanced approach. Modulation of AVP or its receptor pathways, particularly V_1b_ receptor-dependent signaling, could in principle fine-tune pancreatic islet responses under selected conditions. However, inappropriate activation or desensitization of α and β cells by fluctuating or persistently elevated AVP may also worsen glycemic instability, especially in patients experiencing frequent transitions between hyperglycemia and hypoglycemia.

In summary, AVP plays an important role in modulating pancreatic endocrine function, but its effects depend strongly on glucose concentration, cAMP signaling, and the current dynamic state of the islet. The difficulty of maintaining AVP within a physiological range during hyperosmolality, dehydration, or hypoglycemia underscores the need to consider AVP signaling in the broader neurohormonal regulation of diabetes, particularly in relation to insulin secretion, glucagon counterregulation, and prevention of hypoglycemic episodes.

This study has several limitations. Although pancreatic slices preserve local tissue architecture and enable high-resolution analysis of α and β cell Ca²⁺ dynamics, they do not fully reproduce the intact in vivo environment, including vascular perfusion, innervation, and systemic hormonal regulation. Ca²⁺ imaging is a surrogate readout for hormone secretion; although we expanded insulin and glucagon secretion measurements, the used stimulation protocols could not establish a strict quantitative relationship between Ca²⁺ dynamics and secretory output. In addition, most experiments were performed under cAMP-permissive conditions, which increases sensitivity for detecting AVP-dependent modulation but complicates the separation of direct beta cell effects from intra-islet interactions. Finally, we were not yet able to sufficiently standardize or dynamically monitor the metabolic state of individual islets, which remains an important source of variability and a key objective for future studies.

## Acknowledgements

MSR received grants by the Austrian Science Fund/Fonds zur Förderung der Wissenschaftlichen Forschung (bilateral grants I3562-B27 and I4319-B30), a grant from Vienna Science and Technology Fund WWTF (LS23-026), and financial support from NIH (R01DK127236). MSR, AS and DK further received financial support from the Slovenian Research Agency (research core funding program P3-0396). Research in the laboratory of CWG has been supported by a grant from the Austrian Science Fund (FWF) with grant-DOI: 10.55776/P36762. Research in MM’s laboratory has been funded by the European Research Council (ERC) under the European Union’s Horizon 2020 research and innovation program (714366) and the Australian Research Council (ARC, FT210100266). M.P. has been supported by the Austrian Academy of Sciences (ÖAW) through a DOC fellowship (27012).

## Author Contributions

K., N. M., L. K. B., E. P. L, J. P., S. P. performed pancreatic tissue slice imaging experiments. N. M. and J. P. performed hormone release assays, S. P. performed RNAscope experiments. M. S. R and J. P. analyzed the data and created the figures. M. S. R., J. K., L. K. B., C. W. G, drafted the paper and all authors edited the paper. C. W. G,, M. P., M. M., and X. K. provided resources. M. S. R., A. S., D. K., C. W. G., and M. M. acquired funding.

## Conflict of interest

The authors declare no competing interests.

## Data availability

Raw data supporting the peptide synthesis & analysis, as well as pharmacological and physiological measurements of this study are available from the corresponding authors upon reasonable request.

---

**Supplementary Figure 1:**
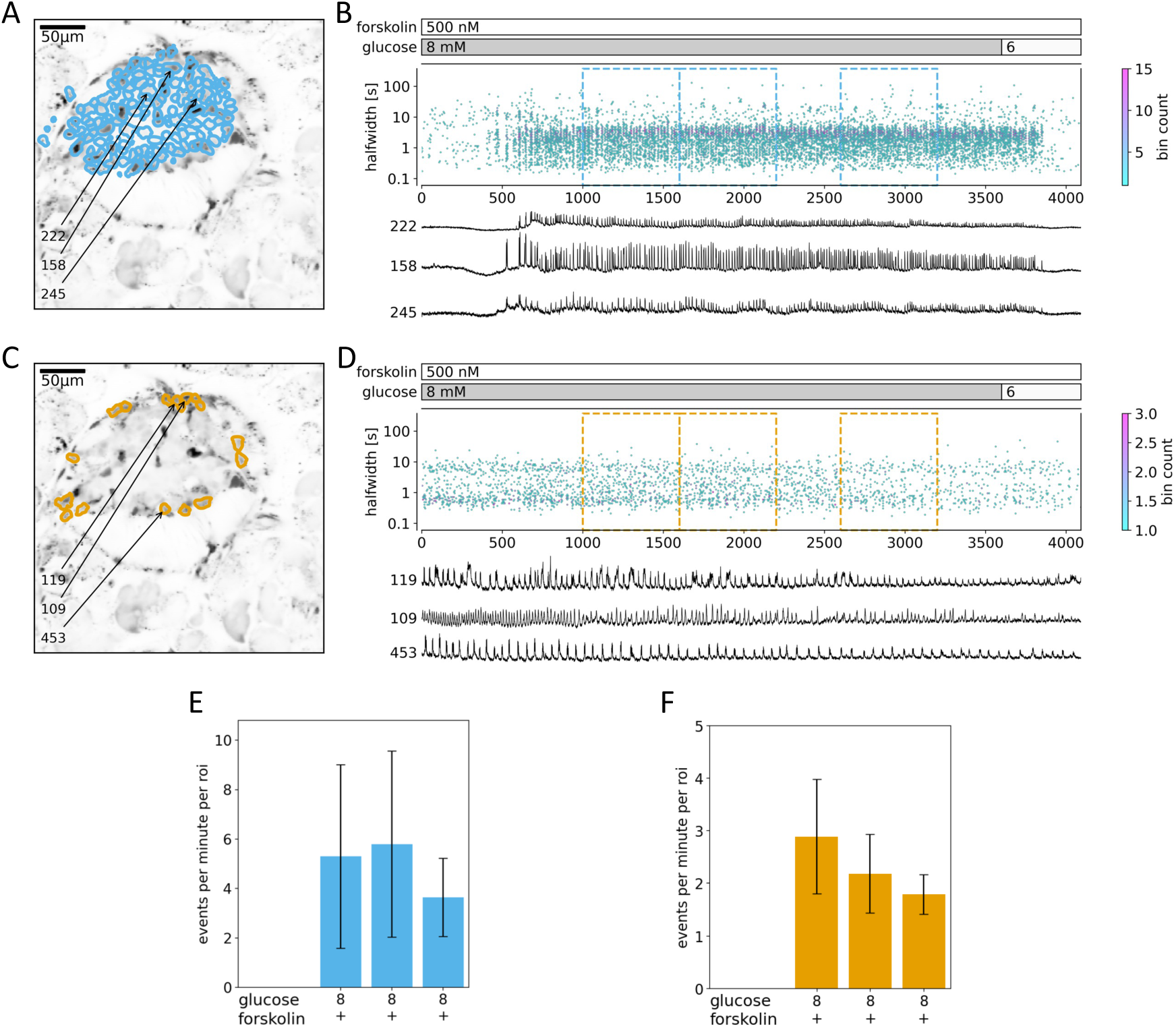
The effect of physiological levels of glucose and forskolin on β (A, B, E, G, H) and α (C, D, F) cells. (A, C) Indicated are the ROIs whose filtered traces correlate best with the average trace for the whole islet. (B, D) (top panel) Time distribution of all measured eventś halfwidths at their peak times. Stimulation protocol is indicated above. The color bar indicates the bin count in each time/halfwidth point. The dashed-line rectangles indicate the 3 time periods where events per minute per ROI parameter was sampled for panels E and F. (bottom panel) Representative traces from ROIs indicated in A and C. (E, F) Prolonged exposure to physiological glucose stimulation and low forskolin concentration resulted in a stable Ca^2+^ oscillations over almost an hour in β and α cells. The parameters were distributed normally, and we used one-way ANOVA and Tukey post-hoc test. The data are presented as mean +/− SD.

**Supplementary Figure 2:**
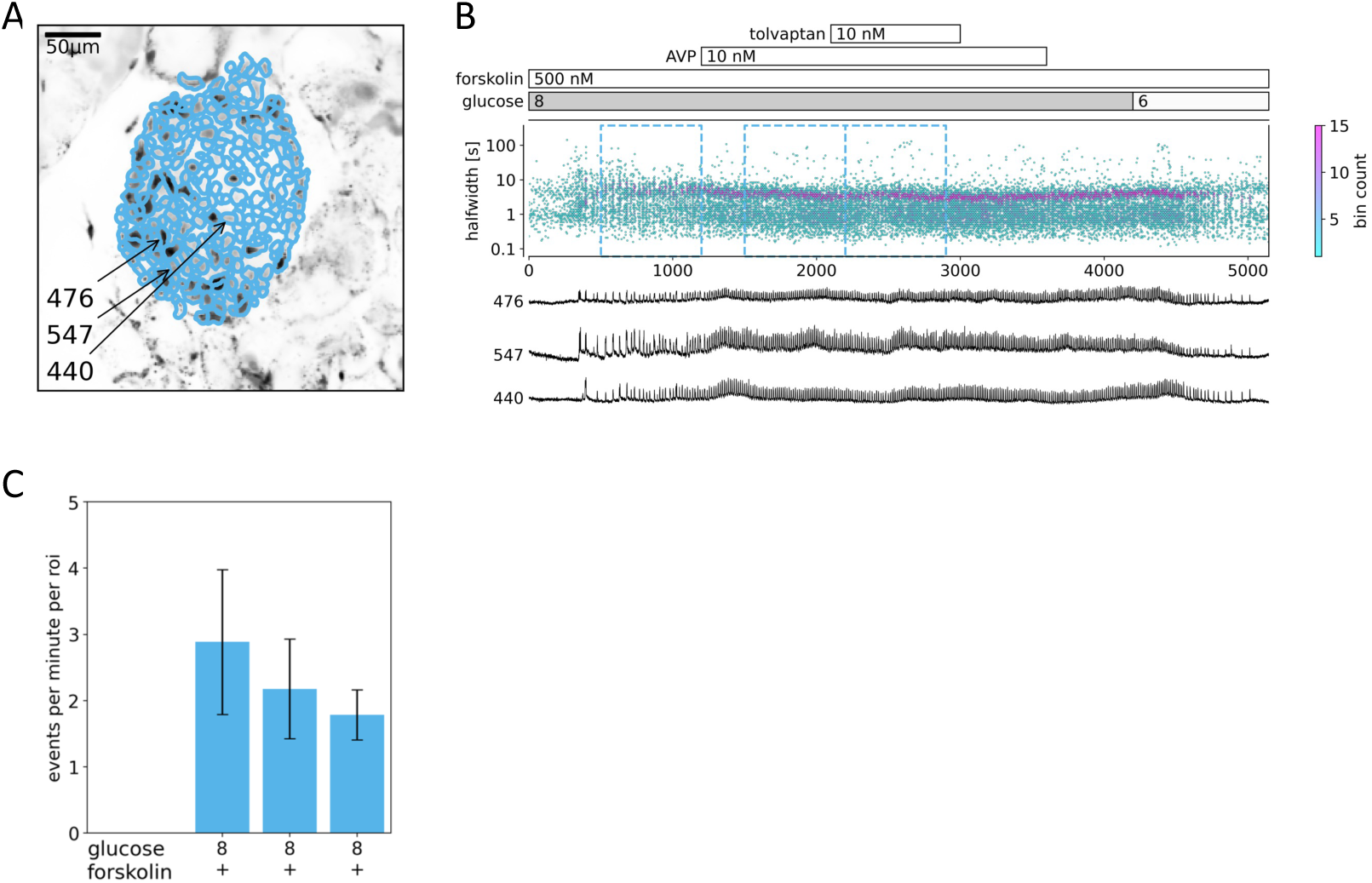
Tolvaptan, a V_2_ receptor antagonist, has no significant effect on β cells. (A) Indicated are the ROIs whose filtered traces correlate best with the average trace for the whole islet. (B) (top panel) Time distribution of all measured eventś halfwidths at their peak times. Stimulation protocol is indicated above. The color bar indicates the bin count in each time/halfwidth point. The dashed-line rectangles indicate the 3 time periods where events per minute per ROI parameter was sampled for the panel. (bottom panel) Representative traces from ROIs indicated in A (C) Tolvaptan had no significant effects on the number of events per minute per ROI. The parameters were distributed normally, and we used one-way ANOVA and Tukey post-hoc test. The data are presented as mean +/− SD.

**Supplementary Figure 3:**
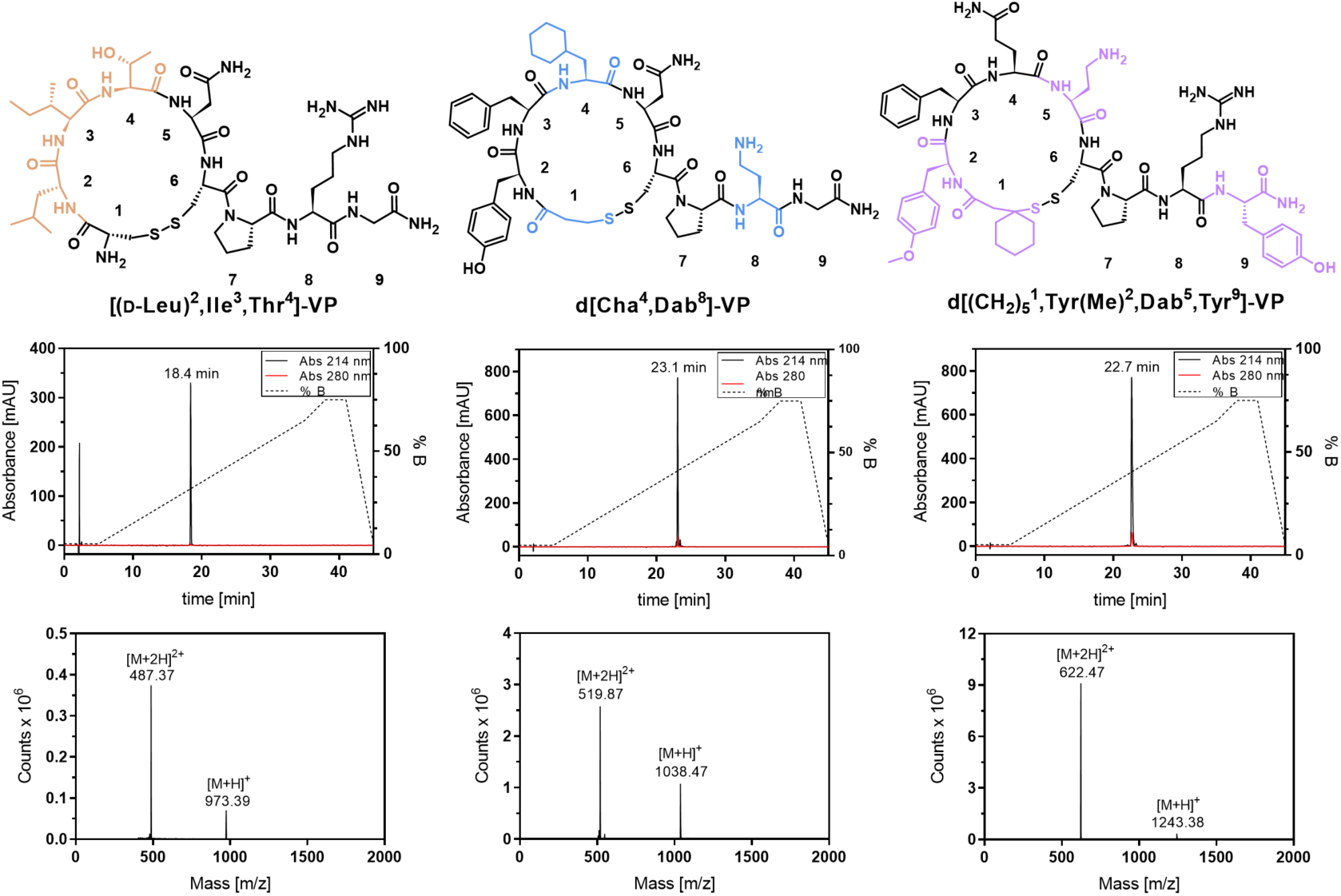
Overview of molecular structures, chromatograms, and mass spectra of synthesized compounds. Amino acid residues that differ from native vasopressin were marked in different colors. Analytic RP-HPLC chromatograms were measured for product purity determination at 214 nm. Solvent A (ddH_2_O + 0.1% TFA) and B (ACN + 0.08% TFA) were used as eluents at 1 mL/min flow rate and a linear gradient of 5-65% B in 30 min on a Thermo Fisher Scientific Vanquish Horizon UHPLC system using a Kromasil Classic C_18_ column (4.6 x 150 mm, 300 Å, 5 µm). Mass spectra were measured for product identity verification through direct injection on a Thermo Scientific Dionex Ultimate 3000 system equipped with a Thermo Scientific MSQ Plus ESI-MS unit (positive ionization mode).

**Supplementary Table 1:**
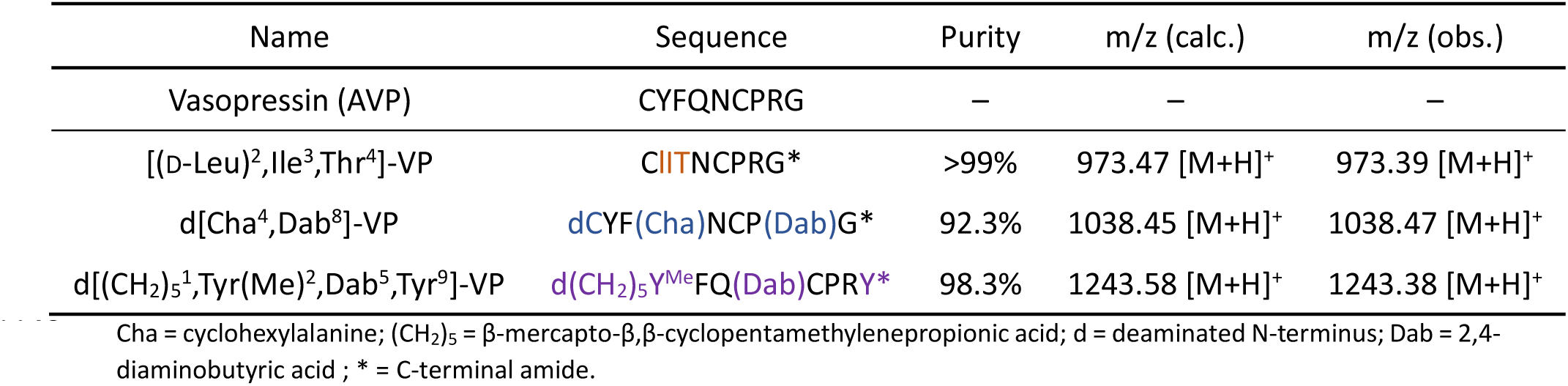
Name, sequence, purity, and mass identity of synthesized vasopressin analogs and vasopressin. Amino acid residues that differ from native vasopressin were marked in different colors.

| Name | Sequence | Purity | m/z (calc.) | m/z (obs.) |
| --- | --- | --- | --- | --- |
| Vasopressin (AVP) | CYFQNCPRG | — | — | — |
| [(D-Leu)<sup>2</sup>,Ile<sup>3</sup>,Thr<sup>4</sup>]-VP | CIITNCPRG* | >99% | 973.47 [M+H]<sup>+</sup> | 973.39 [M+H]<sup>+</sup> |
| d[Cha<sup>4</sup>,Dab<sup>8</sup>]-VP | dCYF(Cha)NCP(Dab)G* | 92.3% | 1038.45 [M+H]<sup>+</sup> | 1038.47 [M+H]<sup>+</sup> |
| d[(CH<sub>2</sub>)<sub>5</sub><sup>1</sup>,Tyr(Me)<sup>2</sup>,Dab<sup>5</sup>,Tyr<sup>9</sup>]-VP | d(CH<sub>2</sub>)<sub>5</sub>Y<sup>Me</sup>FQ(Dab)CPRY* | 98.3% | 1243.58 [M+H]<sup>+</sup> | 1243.38 [M+H]<sup>+</sup> |

Cha = cyclohexylalanine; (CH<sub>2</sub>)<sub>5</sub> = β-mercapto- $\beta$ , β-cyclopentamethylenepropionic acid; d = deaminated N-terminus; Dab = 2,4-diaminobutyric acid ; \* = C-terminal amide.

## Notes

### Competing Interest Statement

The authors have declared no competing interest.

### Summary of Updates

Change in the title. Added experiment and a major revision after peer review.

## References

1. Bankir L. Antidiuretic action of vasopressin: quantitative aspects and interaction between V1a and V2 receptor-mediated effects. Cardiovasc Res. 2001;51(3):372–90.

2. Pittman QJ. Vasopressin and central control of the cardiovascular system: A 40-year retrospective. J Neuroendocrinol. 2021;33(11):e13011.

3. Mavani GP, DeVita MV, Michelis MF. A review of the nonpressor and nonantidiuretic actions of the hormone vasopressin. Front Med (Lausanne). 2015;2:19.

4. Yoshimura M, Conway-Campbell B, Ueta Y. Arginine vasopressin: Direct and indirect action on metabolism. Peptides. 2021;142:170555.

5. Hems DA, Whitton PD. Stimulation by vasopressin of glycogen breakdown and gluconeogenesis in the perfused rat liver. Biochem J. 1973;136(3):705–9.

6. Dunning BE, Moltz JH, Fawcett CP. Actions of neurohypophysial peptides on pancreatic hormone release. Am J Physiol. 1984;246(1 Pt 1):E108–14.

7. Oshikawa S, Tanoue A, Koshimizu TA, Kitagawa Y, Tsujimoto G. Vasopressin stimulates insulin release from islet cells through V1b receptors: a combined pharmacological/knockout approach. Mol Pharmacol. 2004;65(3):623–9.

8. Mohan S, Moffett RC, Thomas KG, Irwin N, Flatt PR. Vasopressin receptors in islets enhance glucose tolerance, pancreatic beta-cell secretory function, proliferation and survival. Biochimie. 2019;158:191–8.

9. Fujiwara Y, Hiroyama M, Sanbe A, Yamauchi J, Tsujimoto G, Tanoue A. Mutual regulation of vasopressin- and oxytocin-induced glucagon secretion in V1b vasopressin receptor knockout mice. J Endocrinol. 2007;192(2):361–9.

10. Folny V, Raufaste D, Lukovic L, Pouzet B, Rochard P, Pascal M, et al. Pancreatic vasopressin V1b receptors: characterization in In-R1-G9 cells and localization in human pancreas. Am J Physiol Endocrinol Metab. 2003;285(3):E566–76.

11. Korošak D, Postić S, Stožer A, Rupnik MS. Collective biological computation in metabolic economy. 4open. 2023;6:3.

12. Hawthorn J, Ang VT, Jenkins JS. Localization of vasopressin in the rat brain. Brain Res. 1980;197(1):75–81.

13. Kim A, Knudsen JG, Madara JC, Benrick A, Hill TG, Abdul Kadir L, et al. Arginine-vasopressin mediates counter-regulatory glucagon release and is diminished in type 1 diabetes. Elife. 2021;10.

14. Dunn FL, Brennan TJ, Nelson AE, Robertson GL. The role of blood osmolality and volume in regulating vasopressin secretion in the rat. J Clin Invest. 1973;52(12):3212–9.

15. Beardwell CG, Geelen G, Palmer HM, Roberts D, Salamonson L. Radioimmunoassay of plasma vasopressin in physiological and pathological states in man. J Endocrinol. 1975;67(2):189–202.

16. Roberts EM, Newson MJ, Pope GR, Landgraf R, Lolait SJ, O’Carroll AM. Abnormal fluid homeostasis in apelin receptor knockout mice. J Endocrinol. 2009;202(3):453–62.

17. Mechaly I, Macari F, Laliberte MF, Lautier C, Serrano JJ, Cros G, et al. Identification by RT-PCR and immunolocalization of arginine vasopressin in rat pancreas. Diabetes Metab. 1999;25(6):498–501.

18. Abu-Basha EA, Yibchok-Anun S, Hsu WH. Glucose dependency of arginine vasopressin-induced insulin and glucagon release from the perfused rat pancreas. Metabolism. 2002;51(9):1184–90.

19. Berggren PO, Ostenson CG, Petersson B, Hellman B. Evidence for divergent glucose effects on calcium metabolism in pancreatic beta- and alpha 2-cells. Endocrinology. 1979;105(6):1463–8.

20. Gao ZY, Drews G, Nenquin M, Plant TD, Henquin JC. Mechanisms of the stimulation of insulin release by arginine-vasopressin in normal mouse islets. J Biol Chem. 1990;265(26):15724–30.

21. Knudtzon J. Acute effects of oxytocin and vasopressin on plasma levels of glucagon, insulin and glucose in rabbits. Horm Metab Res. 1983;15(2):103–4.

22. Mineo H, Ito M, Muto H, Kamita H, Hyun HS, Onaga T, et al. Effects of oxytocin, arginine-vasopressin and lysine-vasopressin on insulin and glucagon secretion in sheep. Res Vet Sci. 1997;62(2):105–10.

23. Altszuler N, Hampshire J. Oxytocin infusion increases plasma insulin and glucagon levels and glucose production and uptake in the normal dog. Diabetes. 1981;30(2):112–4.

24. Spruce BA, McCulloch AJ, Burd J, Orskov H, Heaton A, Baylis PH, et al. The effect of vasopressin infusion on glucose metabolism in man. Clin Endocrinol (Oxf). 1985;22(4):463–8.

25. Khalaf LJ, Taylor KW. Pertussis toxin reverses the inhibition of insulin secretion caused by [Arg8]vasopressin in rat pancreatic islets. FEBS Lett. 1988;231(1):148–50.

26. Koshimizu TA, Nakamura K, Egashira N, Hiroyama M, Nonoguchi H, Tanoue A. Vasopressin V1a and V1b receptors: from molecules to physiological systems. Physiol Rev. 2012;92(4):1813–64.

27. Robben JH, Knoers NVAM, Deen PMT. Cell biological aspects of the vasopressin type-2 receptor and aquaporin 2 water channel in nephrogenic diabetes insipidus. American Journal of Physiology - Renal Physiology. 2006;291(2):F257–F70.

28. van der Meulen T, Mawla AM, DiGruccio MR, Adams MW, Nies V, Dolleman S, et al. Virgin Beta Cells Persist throughout Life at a Neogenic Niche within Pancreatic Islets. Cell Metab. 2017;25(4):911–26 e6.

29. Chen TH, Lee B, Hsu WH. Arginine vasopressin-stimulated insulin secretion and elevation of intracellular Ca++ concentration in rat insulinoma cells: influences of a phospholipase C inhibitor 1-[6-[[17 beta-methoxyestra-1,3,5(10)-trien-17-yl]amino]hexyl]-1H-pyrrole-2,5-dione (U-73122) and a phospholipase A2 inhibitor N-(p-amylcinnamoyl)anthranilic acid. J Pharmacol Exp Ther. 1994;270(3):900–4.

30. Fujiwara Y, Hiroyama M, Sanbe A, Aoyagi T, Birumachi J, Yamauchi J, et al. Insulin hypersensitivity in mice lacking the V1b vasopressin receptor. J Physiol. 2007;584(Pt 1):235–44.

31. Nakamura K, Yamashita T, Fujiki H, Aoyagi T, Yamauchi J, Mori T, et al. Enhanced glucose tolerance in the Brattleboro rat. Biochem Biophys Res Commun. 2011;405(1):64–7.

32. O’Carroll AM, Howell GM, Roberts EM, Lolait SJ. Vasopressin potentiates corticotropin-releasing hormone-induced insulin release from mouse pancreatic beta-cells. J Endocrinol. 2008;197(2):231–9.

33. Tse A, Lee AK, Tse FW. Role of the TWIK-Related Potassium (TREK)-1 Channels in the Regulation of Adrenocorticotropic Hormone (ACTH) Secretion from Pituitary Corticotropes. Masterclass in Neuroendocrinology. 8 2020. p. 219–39.

34. Kjaer A, Knigge U, Bach FW, Warberg J. Permissive, mediating and potentiating effects of vasopressin in the ACTH and beta-endorphin response to histamine and restraint stress. Neuroendocrinology. 1993;58(5):588–96.

35. Leech CA, Dzhura I, Chepurny OG, Kang G, Schwede F, Genieser H-G, et al. Molecular physiology of glucagon-like peptide-1 insulin secretagogue action in pancreatic β cells. Progress in Biophysics and Molecular Biology. 2011;107(2):236–47.

36. Quesada I, Tudurí E, Ripoll C, Nadal Á. Physiology of the pancreatic α-cell and glucagon secretion: role in glucose homeostasis and diabetes. Journal of Endocrinology. 2008;199(1):5–19.

37. Bezprozvanny l, Watras J, Ehrlich BE. Bell-shaped calcium-response curves of lns(l,4,5)P3- and calcium-gated channels from endoplasmic reticulum of cerebellum. Nature. 1991;351(6329):751–4.

38. Stozer A, Dolensek J, Krizancic Bombek L, Pohorec V, Slak Rupnik M, Klemen MS. Confocal Laser Scanning Microscopy of Calcium Dynamics in Acute Mouse Pancreatic Tissue Slices. J Vis Exp. 2021(170).

39. Wang F, Flanagan J, Su N, Wang LC, Bui S, Nielson A, et al. RNAscope: a novel in situ RNA analysis platform for formalin-fixed, paraffin-embedded tissues. J Mol Diagn. 2012;14(1):22–9.

40. Kvalseth TO. Regression models of emergency medical service demand for different types of emergencies. IEEE Trans Syst Man Cybern. 1979;9(1):10–7.

41. Soille PJ, Ansoult MM. Automated basin delineation from digital elevation models using mathematical morphology. Signal Processing. 1990;20(2):171–82.

42. Lindeberg T. Scale-space theory in computer vision: Springer Science & Business Media; 2013.

43. Kremsmayr T, Muttenthaler M. Fmoc Solid Phase Peptide Synthesis of Oxytocin and Analogues. Methods Mol Biol. 2022;2384:175–99.

44. Kremsmayr T, Aljnabi A, Blanco-Canosa JB, Tran HNT, Emidio NB, Muttenthaler M. On the Utility of Chemical Strategies to Improve Peptide Gut Stability. J Med Chem. 2022;65(8):6191–206.

45. Koehbach J, O’Brien M, Muttenthaler M, Miazzo M, Akcan M, Elliott AG, et al. Oxytocic plant cyclotides as templates for peptide G protein-coupled receptor ligand design. Proc Natl Acad Sci U S A. 2013;110(52):21183–8.

46. Duerrauer L, Muratspahic E, Gattringer J, Keov P, Mendel HC, Pfleger KDG, et al. I8-arachnotocin-an arthropod-derived G protein-biased ligand of the human vasopressin V(2) receptor. Sci Rep. 2019;9(1):19295.

47. Di Giglio MG, Muttenthaler M, Harpsoe K, Liutkeviciute Z, Keov P, Eder T, et al. Development of a human vasopressin V(1a)-receptor antagonist from an evolutionary-related insect neuropeptide. Sci Rep. 2017;7:41002.

48. Cheng Y, Prusoff WH. Relationship between the inhibition constant (K1) and the concentration of inhibitor which causes 50 per cent inhibition (I50) of an enzymatic reaction. Biochem Pharmacol. 1973;22(23):3099–108.

49. Postic S, Sarikas S, Pfabe J, Pohorec V, Krizancic Bombek L, Sluga N, et al. High-resolution analysis of the cytosolic Ca(2+) events in beta cell collectives in situ. Am J Physiol Endocrinol Metab. 2023;324(1):E42–E55.

50. Postic S, Pfabe J, Sarikas S, Ehall B, Pieber T, Korosak D, et al. Tracking Ca2+ Dynamics in NOD Mouse Islets During Spontaneous Diabetes Development. Diabetes. 2023;72(9):1251–61.

51. Sluga N, Križančić Bombek L, Kerčmar J, Sarikas S, Postić S, Pfabe J, et al. Physiological levels of adrenaline fail to stop pancreatic beta cell activity at unphysiologically high glucose levels. Front Endocrinol (Lausanne). 2022;13:1013697.

52. Sanchez-Andrés JV, Soria B. Muscarinic inhibition of pancreatic B-cells. European Journal of Pharmacology. 1991;205(1):89–91.

53. Pena A, Murat B, Trueba M, Ventura MA, Wo NC, Szeto HH, et al. Design and synthesis of the first selective agonists for the rat vasopressin V(1b) receptor: based on modifications of deamino-[Cys1]arginine vasopressin at positions 4 and 8. J Med Chem. 2007;50(4):835–47.

54. Chan WY, Wo NC, Cheng LL, Manning M. Isosteric substitution of Asn5 in antagonists of oxytocin and vasopressin leads to highly selective and potent oxytocin and V1a receptor antagonists: new approaches for the design of potential tocolytics for preterm labor. J Pharmacol Exp Ther. 1996;277(2):999–1003.

55. Manning M, Stoev S, Chini B, Durroux T, Mouillac B, Guillon G. Peptide and non-peptide agonists and antagonists for the vasopressin and oxytocin V1a, V1b, V2 and OT receptors: research tools and potential therapeutic agents. Prog Brain Res. 2008;170:473–512.

56. Sanchez-Andres JV, Gomis A, Valdeolmillos M. The electrical activity of mouse pancreatic beta-cells recorded in vivo shows glucose-dependent oscillations. J Physiol. 1995;486 (Pt 1)(Pt 1):223–8.

57. Paradiz Leitgeb E, Pohorec V, Krizancic Bombek L, Skelin Klemen M, Duh M, Gosak M, et al. Calcium Imaging and Analysis in Beta Cells in Acute Mouse Pancreas Tissue Slices. Methods Mol Biol. 2025;2861:223–46.

58. Yun Y, Guo S, Xie X. V1bR enhances glucose-stimulated insulin secretion by paracrine production of glucagon which activates GLP-1 receptor. Cell Biosci. 2024;14(1):110.

59. van Krieken PP, Voznesenskaya A, Dicker A, Xiong Y, Park JH, Lee JI, et al. Translational assessment of a genetic engineering methodology to improve islet function for transplantation. EBioMedicine. 2019;45:529–41.

60. Ren H, Li Y, Xie B, Fu Z, Peng X, Qian W, et al. Pancreatic islet oscillation rhythmicity arises from delta and alpha cell interactions. Cell Syst. 2026:101587.

61. Marciniak A, Cohrs CM, Tsata V, Chouinard JA, Selck C, Stertmann J, et al. Using pancreas tissue slices for in situ studies of islet of Langerhans and acinar cell biology. Nat Protoc. 2014;9(12):2809–22.

62. Orcel H, Albizu L, Perkovska S, Durroux T, Mendre C, Ansanay H, et al. Differential coupling of the vasopressin V1b receptor through compartmentalization within the plasma membrane. Mol Pharmacol. 2009;75(3):637–47.

63. Taylor CW, Tovey SC. IP(3) receptors: toward understanding their activation. Cold Spring Harb Perspect Biol. 2010;2(12):a004010.

64. Koceva A, Janez A, Jensterle M. The Impact of Hydration on Metabolic Outcomes: From Arginine-Vasopressin Signaling to Clinical Implications. Medicina (Kaunas). 2025;61(5).

65. Politi A, Gaspers LD, Thomas AP, Hofer T. Models of IP3 and Ca2+ oscillations: frequency encoding and identification of underlying feedbacks. Biophys J. 2006;90(9):3120–33.

66. Gilon P, Nenquin M, Henquin JC. Muscarinic stimulation exerts both stimulatory and inhibitory effects on the concentration of cytoplasmic Ca2+ in the electrically excitable pancreatic B-cell. Biochem J. 1995;311 (Pt 1)(Pt 1):259–67.

67. Saez M, Briscoe J, Rand DA. Dynamical landscapes of cell fate decisions. Interface Focus. 2022;12(4):20220002.

68. Zmazek J, Klemen MS, Markovic R, Dolensek J, Marhl M, Stozer A, et al. Assessing Different Temporal Scales of Calcium Dynamics in Networks of Beta Cell Populations. Front Physiol. 2021;12:612233.

69. Derick S, Cheng LL, Voirol MJ, Stoev S, Giacomini M, Wo NC, et al. [1-deamino-4-cyclohexylalanine] arginine vasopressin: a potent and specific agonist for vasopressin V1b receptors. Endocrinology. 2002;143(12):4655–64.

